# CvkR, a novel MerR-type transcriptional regulator, is a repressor of class 2 type V-K CRISPR-associated transposase systems

**DOI:** 10.1101/2022.05.13.491168

**Authors:** Marcus Ziemann, Viktoria Reimann, Yajing Liang, Yue Shi, Yuman Xie, Hui Li, Tao Zhu, Xuefeng Lu, Wolfgang R. Hess

**Affiliations:** University of Freiburg, Faculty of Biology, Institute of Biology III, Genetics and Experimental Bioinformatics, Schänzlestr. 1, D-79104, Germany; Qingdao Institute of Bioenergy and Bioprocess Technology (QIBEBT), Chinese Academy of Sciences, No.189 Songling Road, Qingdao 266101, China; Shandong Energy Institute, Qingdao 266101, China; Qingdao New Energy Shandong Laboratory, Qingdao 266101, China; University of Chinese Academy of Sciences, Beijing 100049, China; Laboratory for Marine Biology and Biotechnology, Qingdao National Laboratory for Marine Science and Technology, Qingdao 266237, China

**Keywords:** CRISPR, CRISPR-associated transposons, cyanobacteria, transcriptional regulator

## Abstract

CRISPR-associated transposons (CASTs) exist in different groups of bacteria, including certain cyanobacteria, which contain type V-K CAST systems. These systems contain genes encoding Tn7-like transposase subunits and a divergent number of cargo genes. How the activity of these systems is controlled *in situ* has remained largely unknown but possibly regulatory genes within these elements are prime candidates. Deletion of the respective regulator gene *alr3614* in the cyanobacterium *Anabaena (Nostoc)* sp. PCC 7120 led to the overexpression of CRISPR tracrRNA, precursor crRNAs and mRNAs encoding the Cas12k effector protein (*all3613*) and Tn7-like transposase subunits. Upon complementation, these same genes were repressed again. DNase I footprinting and electrophoretic mobility shift assays verified the direct interaction between Alr3614 and the promoter of *cas12k* and identified a widely conserved binding motif. Structural analysis of Alr3614 at 1.5 Å resolution revealed that it belongs to the MerR-type transcription factor family but with distinct dimerization and effector-binding domains. This protein assembles into a homodimer interacting with DNA through its N-terminal winged helix-turn-helix (wHTH) domain and binds an effector molecule through a C-terminal α-helical domain lacking a conserved cysteine. These results identify Alr3614 as a transcriptional repressor of the CAST system in *Anabaena* sp. PCC 7120. We suggest naming this family of repressors CvkR for Cas V-K repressors, which are at the core of a widely conserved regulatory mechanism that controls type V-K CAST systems.

## Introduction

Native Clustered Regularly Interspaced Short Palindromic Repeats (CRISPRs) and CRISPR-associated (Cas) proteins are well characterized for their function as RNA-based adaptive and inheritable immune systems found in many bacteria and archaea^1–6^. Multiple genetic approaches developed from these native CRISPR-Cas systems have become popular for the manipulation of gene expression and genome editing^7–9^. CRISPR-Cas systems are extremely diverse and are classified into 2 classes, 6 types and 33 subtypes^10^. Recently, a remarkable group of derivatives has been discovered that constitute hybrids of Tn7-like transposons and CRISPR systems encoding Cas12k effectors with naturally inactivated nuclease domains^11^ or encoding Cascade complexes lacking the Cas3 nuclease component^12, 13^. The respective transposon-associated CRISPR systems include class 1 type I-F, I-B, and class 2 type V-K systems^13^. These systems, called CRISPR-associated transposons (CASTs), are capable of catalyzing the transposition of mobile genetic elements guided by crRNAs, while the type I-B associated systems use a dedicated TniQ/TnsD protein-based homing mechanism^13^.

The system characterized from *Vibrio cholerae* consists of genes encoding the subtype I-F CRISPR-Cas proteins Cas6, Cas7, and Cas8 and genes encoding the transposon proteins TnsA, TnsB, TnsC and TniQ^12^. A single instance of a class 1 type I-B and of several class 2 type V-K CAST systems have been reported in several different cyanobacteria, which constitute a rich natural resource for these systems^11, 13, 14^.

The V-K CAST systems, first characterized in *Scytonema hofmanni*^11^, contain genes encoding the effector complex subunit Cas12k and the Tn7-like transposase subunits TnsB, TnsC and TniQ, while *tnsA* is lacking. Targeting transposition by these CAST systems depends on the DNA-crRNA interaction facilitated by the effector protein Cas12k^11^. The TnsC transposon then forms helical polymers around the DNA supported by ATP binding^15^. The growth in the 5’ to 3’ direction is stopped by TniQ binding at the filament end concomitantly connecting the TnsC filament with Cas12k. On the other filament end, the Mu-like transposase TnsB then starts to integrate the transposon^15^. In addition to the genes encoding transposase and effector proteins, all of these systems contain various numbers of cargo genes. Novel genetic approaches have been developed from the different Tn7-CRISPR–Cas hybrid systems^16–18^, underlining that the better characterization of such systems is of both fundamental and applied interest.

While the paradigm is that native CRISPR-Cas systems primarily protect genome integrity against mobile genetic elements, the CAST systems seem to violate this paradigm since they constitute transposable elements by definition. Thus, the tight regulation of these systems can be expected. Indeed, CAST systems have also been reported to contain a gene encoding a putative MerR-type transcriptional regulator^14^, but this association has not been systematically investigated, nor has its function been addressed experimentally thus far.

We have studied the CRISPR-Cas systems in *Anabaena* (*Nostoc*) sp. PCC 7120 (from here: *Anabaena* 7120), a multicellular nitrogen-fixing model cyanobacterium with a CRISPR-rich chromosome of eleven CRISPR-like repeat-spacer cassettes. All of them are transcribed^14^, and based on the specificities of the cognate Cas6 maturation endonucleases, five of these arrays were assigned to a type III-D and another five to a type I-D CRISPR-Cas system^19^, while the remaining array (CR_9) belongs to a separate CRISPR type with all the hallmarks of a CAST system^14, 19^.

Here, we first scrutinized the association between putative transcriptional regulators and cyanobacterial CAST systems and found these to belong to four different classes. We then investigated the Alr3614 transcriptional regulator belonging to the *Anabaena* 7120 CRISPR-associated transposase (AnCAST) system. We found that both the *cas12k* gene *all3613* and the *merR*-like gene *alr3614* are translated from leaderless mRNAs. In the deletion mutant Δ*alr3614,* we observed overexpression of the AnCAST core module, while repression was restored if Alr3614 was expressed from a complementing plasmid vector. Hence, Alr3614 functions as a repressor of AnCAST. The crystal structure analysis at 1.5 Å resolution was consistent with the assignment of Alr3614 to the MerR-type family of transcription factors but also revealed specific features within the dimerization and effector-binding domains. It assembles into a homodimer interacting with DNA through its N-terminal winged helix-turn-helix (wHTH) domain and binds effector molecule(s) through its C-terminal domain with a standard α-helix and lacking the cysteine otherwise widely conserved in this type of regulator. We suggest naming Alr3614 and its functional homologs Cas V-K repressor (CvkR) encoded by the gene *cvkR*.

While almost all CAST systems contain a regulatory gene, phylogenetic analyses suggested that different repressor types can be encoded at the corresponding location, belonging to the Arc repressor superfamily (CopG- and Omega-like repressors) and the MerR family, or appearing as a different class of putative HTH domain-containing proteins. Our results illuminate the role of CvkR regulators in controlling the activity of CAST systems.

## Results

### Architecture of cyanobacterial CAST systems

Starting from the known CAST components, we searched for conserved genes and genetic elements in their vicinity. These elements included the left and right ends (LE and RE) of the transposon, the neighboring tRNA, CRISPR arrays and tracrRNA, the transposase genes (*tniQ, tnsC* and *tnsB*), and genes in reverse orientation next to the start codon of *cas12k* predicted to encode small DNA-binding proteins. We identified 118 CAST systems with a clear *cas12k* gene in 88 different strains. The majority of these were found in the Nostocales (60%), Chroococcales (15%), and Pseudanabaenales (11%), complemented by a small number of CAST systems in the Oscillatoriales, Pleurocapsales, Spirulinales and Synechococcales cyanobacteria (**Table S1**). Three additional CAST systems were found in unclassified filamentous cyanobacteria (CCT1, CCP 2 and 4). From this analysis, we could delineate the general structure of this type of CAST system (**Fig. 1**) consistent with previous analyses and extend them^10, 11, 14^.

**Fig. 1.**
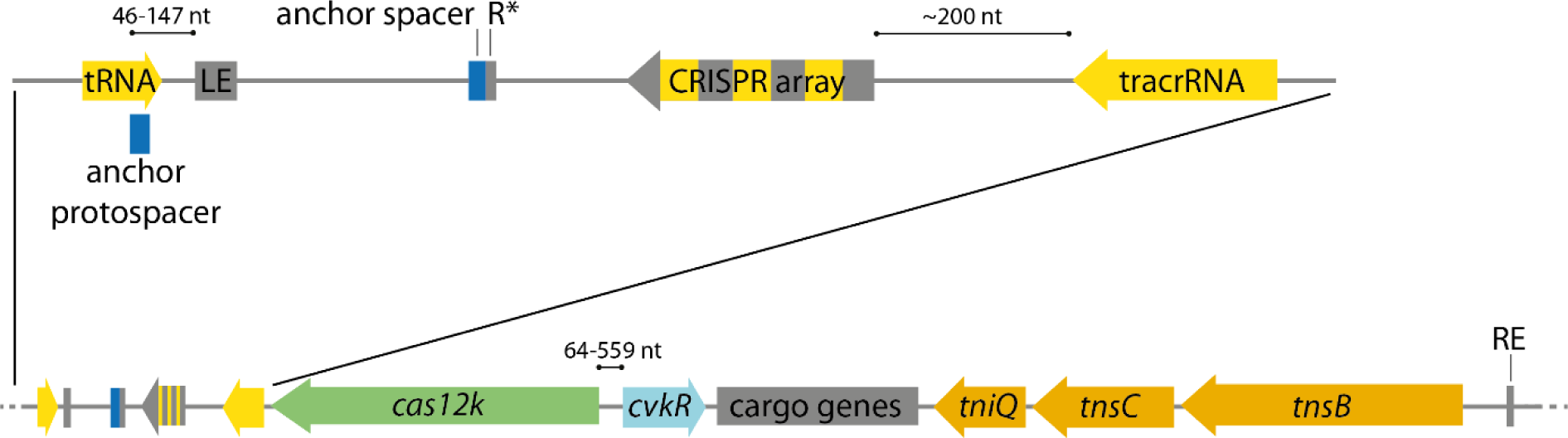
Principal gene arrangement within cyanobacterial CAST systems. The CAST transposon is displayed from its left end (LE) to its right end (RE). The genes are colored according to function (green: *cas12k*, orange: transposase genes, light blue: CAST regulator gene (*cvkR*), yellow: regions from which non-coding RNA is transcribed, dark gray: cargo genes). On top, the region from the tRNA gene to the tracrRNA is magnified. The CRISPR array is depicted with its repeats (gray) and spacers (yellow) separated from the 17 nt anchor spacer (blue) downstream of the array next to a truncated repeat sequence (R*; ∼12 nt). The scheme is not drawn to scale, but distances of particular interest or mentioned in the text are indicated.

The LE usually lies downstream of a tRNA gene oriented toward the transposon. The CRISPR array always follows in a short distance and in reverse orientation with regard to the tRNA gene. The majority of CRISPR repeats are 37 nt long with high conservation at the 3’ end. There is also a sequence-conserved promoter upstream of the tracrRNA, which is followed or, in some cases, even overlapped by the CRISPR-Cas effector gene *cas12k*, transcribed in the same direction.

Next to the LE element, inside the transposon lies a truncated, single repeat downstream but usually clearly separated from the CRISPR array. Directly downstream of this repeat, a truncated spacer sequence of usually 17 nt can be identified that corresponds to a protospacer sequence just outside of the transposon next to the LE, usually within the tRNA gene^13^. The truncated single repeat-spacer sequences, read toward the LE, show a conserved upstream GTN-PAM, consistent with the predicted Cas12k PAM^11^. The distance from this PAM to the LE varies from 46 to 82 nt, with one exception of 147 nt. Because of its conserved position at this site and experimental evidence of CAST integration via protospacer recognition^13^, this motif is likely necessary for the insertion of CAST. We therefore suggest the terms anchor protospacer and anchor spacer for these sequences.

Looking from the other side of the transposon, the first genes next to the RE are the three genes encoding transposase subunits TnsB, TnsC, and TniQ, always in this order, facing away from the RE. The majority of cargo genes, located between *cas12k* and *tniQ*, are significantly more divergent. There is, however, one exception, a gene predicted to encode a small DNA-binding protein next to the start codon of *cas12k* in reverse orientation.

### MerR-type, Arc-type and HTH domain-containing transcriptional regulators are associated with the CAST systems of cyanobacteria

The genes encoding potential CvkRs were identified by their position and orientation with regard to the *cas12k* gene. We analyzed the first gene located upstream of *cas12k* in reverse orientation, assigned it to prominent gene families and then performed similarity searches against NCBI’s non-redundant protein database. In total, we identified 94 CAST systems with genes encoding a putative regulator in a conserved position and orientation with regard to the *cas12k* gene. The *cvkR* genes occur once per CAST system; however, in some instances, degenerated *cvkR* duplicate genes exist immediately after the functional genes. Additionally, we found CAST systems with additional candidate genes but further away from *cas12k,* but none of those instances was considered further to avoid potentially misleading information.

The small DNA-binding proteins encoded by the CAST-associated regulatory genes contain either a helix-turn-helix (HTH) domain or a ribbon-helix-helix domain (RHH) for interaction with DNA. The bioinformatic analysis classified them as members of the MerR family (53 CvkRs), omega-like repressors (22 CvkRs), CopG-like repressors (11 CvkRs) or unspecified HTH domain-containing proteins (8 CvkRs). The MerR family proteins, which also include Alr3614 (NCBI accession: BAB75313.1) from our model organism *Anabaena* 7120, range from 139 to 185 amino acids in length and appear to be monophyletic (**Fig. 2**). Four CvkR proteins differed further in representing fusion proteins with an *hsdR*-restriction domain from a DNA-restriction-methylation (RM) system. RM systems are frequent among the CAST cargo genes, but there is no evidence that RM and CAST systems work together at a mechanistic level. The other HTH proteins are shorter (80-108 residues) and appear polyphyletic (**Fig. 2**). This indicates that these systems gather different regulators independent from one another. The shortest CvkRs are CopG-like proteins and Omega-like repressors containing an RHH DNA-binding domain and ranging from 53 to 72 amino acids in size. For instance, the second CvkR candidate in *Anabaena* 7120 encoded by *asl2690* (BAB74389.1) belongs to the CopG-like family (**Fig. 2** and **Table S1**). Proteins in both of these families can form homodimers, which create a joint antiparallel β-sheet as the basis for DNA binding^20^.

**Fig. 2.**
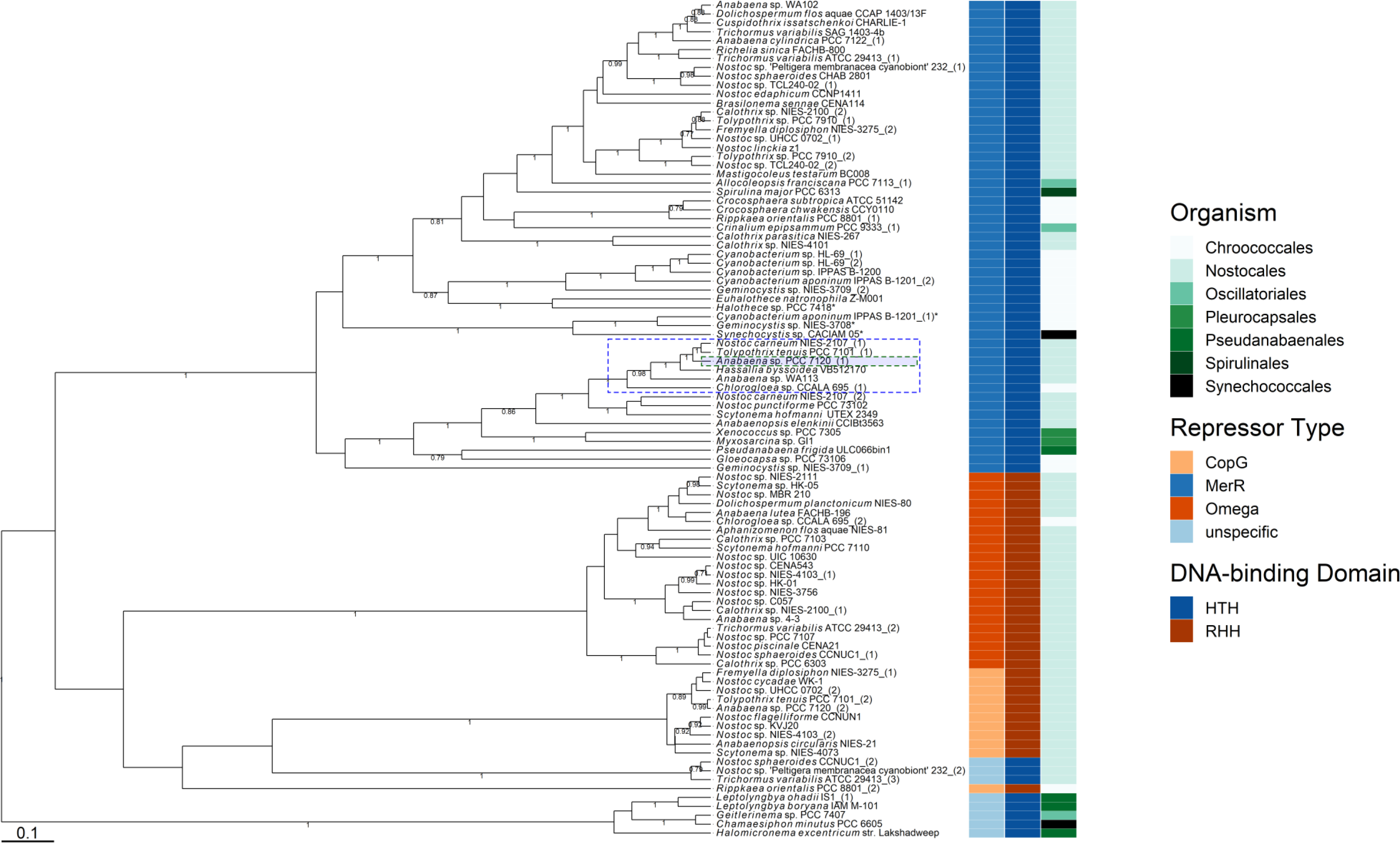
Phylogenetic tree of all CvkR homologs. The identified CvkR proteins were aligned using M-coffee^48, 49^ and analyzed by BEAST^50^. As a prior tree module, we used a Yule process for 1,000,000 states, log processed every 1,000 steps. The resulting tree is depicted with branches labeled with their respective posterior probability until a threshold of 0.5. For better recognition, the proteins were also labeled with their host organism as well as their repressor type and DNA interaction domain. The *Anabaena* 7120 CvkR (Alr3614) (NCBI: BAB75313.1) (green dashed box) is marked as well as its most similar homologs (>70% shared identity, highlighted by a blue-dashed box). Asterisks label four instances of CvkRs fused to an *hsdR* restriction enzyme domain. The multiple sequence alignment is available as **supplemental dataset S1**.

### Expression of Alr3614 from leaderless mRNA

The CvkR encoded by *alr3614* was annotated as a 168 amino acid-long MerR protein. However, sequence comparison of Alr3614 against other MerR-type CvkRs indicated that the NCBI annotation for this protein (here called CvkR-L) was too long (supported by 47 out of 50 homologs), and the actual protein would be 18 residues shorter (here called CvkR-S; **Fig. 3A**). Furthermore, the distance between *cas12k* and genes encoding CopG- and Omega-like CvkRs was longer than that between *cas12k* and genes encoding MerR-type CvkR proteins (usually <100 nt) (**Fig. 3B**).

**Fig. 3.**
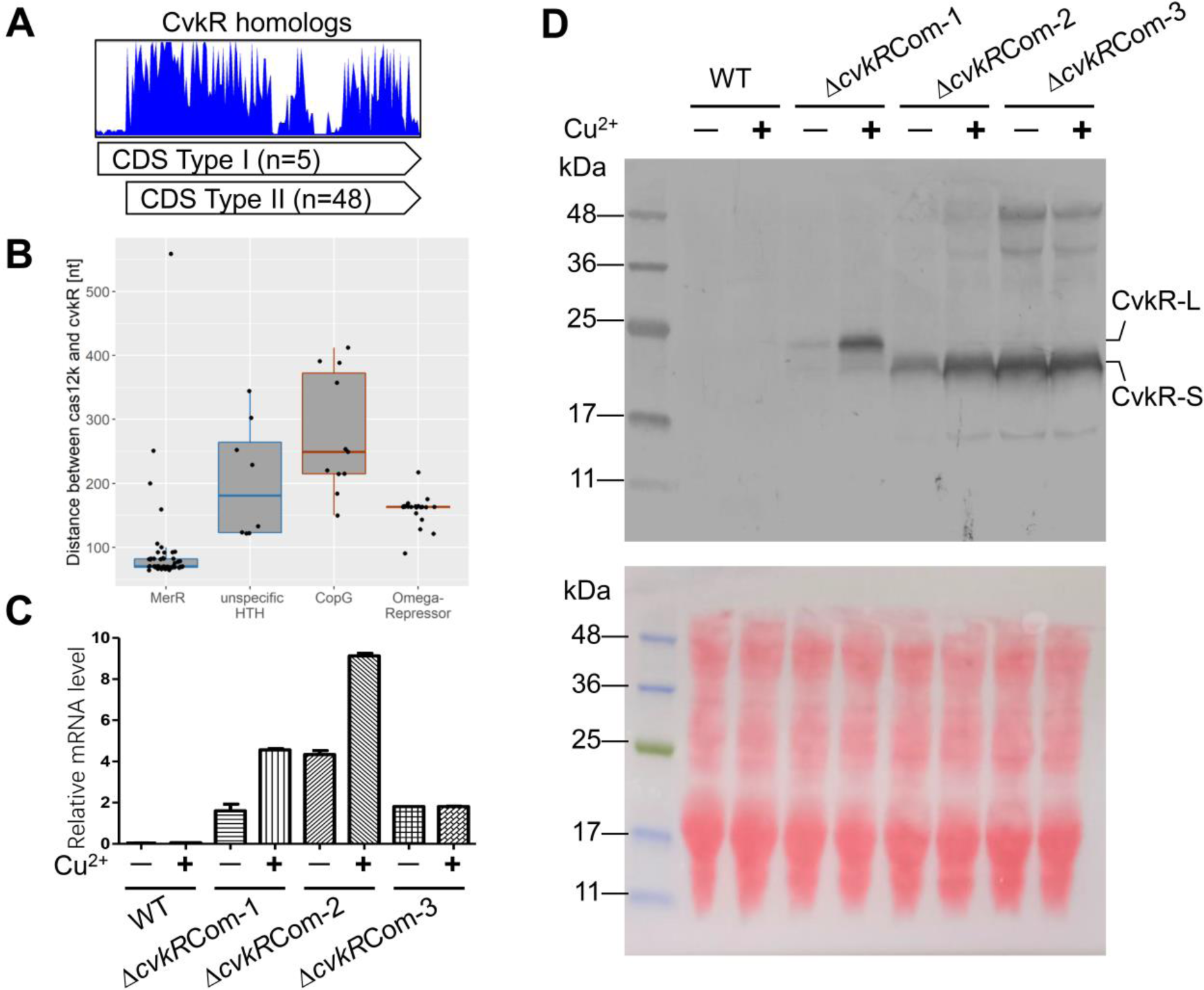
Leaderless expression of *cvkR* genes. **A.** Deduced amino acid sequence alignment of 53 MerR-family CvkR homologs. The sequences of the MerR-family CvkR homologs for the multiple sequence alignment were recovered from public databases and are available as **supplemental dataset S2.** The sequences were aligned using MAFFT^60^ and visualized with UGENE^61^. Only the N-terminal ∼200 amino acids are displayed for clarity reasons. **B.** Distances between the *cas12k* effector complex and *cvkR* genes are plotted according to the respective type of regulator, MerR-like (53 instances), CopG-like (11 cases), Omega-like (22 cases) and nonspecific HTH domain-containing proteins (8 cases). The box plots are colored according to the respective DNA-binding domain (HTH: blue; RHH: red). **C.** qRT–PCR analyses verify that *cvkR* is transcribed in Δ*cvkR*Com strains. The amounts of *cvkR* transcripts were normalized to those of *rnpB* as an internal standard. Three independent experiments were performed, which showed consistent results. **D**. Western blot analyses confirmed the leaderless expression of CvkR. Upper panel, Western blot against the C-terminal 3xFLAG tag; Lower panel, ponceau S staining shows that equal amounts of protein were loaded (100 μg). The calculated molecular masses for CvkR-S and CvkR-L were 20.03 kDa and 22.41 kDa, respectively. Two independent experiments were performed, which showed consistent results. The Δ*cvkR*Com-1, 2 and 3 strains are detailed in **Fig. S1C**.

Moreover, the start codon of *alr3614S* coincides with the previously mapped transcriptional start site (TSS) of its mRNA^21^, suggesting translation of CvkR-S from a leaderless mRNA. The TSS of *cas12k* coincides with the first nucleotide of the start codon as well; therefore, Cas12k is likely also translated from leaderless mRNA. Both genes and their TSSs are separated by an intergenic spacer of 82 nt in *Anabaena* 7120, and their homologs in other species are probably leaderless as well, judged by the generally shorter distance between the two genes than for other types of CvkR-encoding genes (**Fig. 3B**).

To verify the leaderless expression of CvkR-S, we constructed an *alr3614* deletion mutant (Δ*cvkR*) using the CRISPR-Cas12a (Cpf1) genome editing tool (**Fig. S1A and B**) and three versions of complementation mutants (Δ*cvkR*Com-1 to -3). For complementation, shuttle vectors were used, which carried *cvkR* genes driven by the copper-inducible *petE* promoter for the long form CvkR-L (Δ*cvkR*Com-1) or the short form CvkR-S, either containing a leader sequence (Δ*cvkR*Com-3) or not (Δ*cvkR*Com-2). For detection, a 3xFLAG epitope-encoding tag was fused to all three *cvkR* reading frames. The complementing plasmids were introduced into the Δ*cvkR* deletion background and verified (**Fig. S1C and D**). The *cvkR* gene was transcribed in all three constructs (**Fig. 3C**). Translation of the encoded proteins was detected by Western blot, with clear size differences between CvkR-L and CvkR-S (**Fig. 3D**). Intriguingly, in Δ*cvkR*Com-1, we detected not only the CvkR-L form but also a small amount of CvkR-S, suggesting a propensity for initiation of translation at codon 18 even if the mRNA was 5’ elongated. As there was no 5’UTR in Δ*cvkR*Com-1 following P_*petE*_, it seems that the A in the start codon of CvkR-S could also serve as a start site of translation if combined with transcription from a strong promoter. The incongruence between *cvkR* mRNA and protein levels within Δ*cvkR*Com-3 implied that the 5’UTR of P_*petE*_ might impact the transcription rate, mRNA stability and/or translation efficiency of the *cvkR-S* mRNA. Taken together, we provide solid evidence that CvkR could be expressed from leaderless mRNA (Δ*cvkR*Com-2) or the start codon corresponding to codon 18 of the original annotation (Δ*cvkR*Com-3). Because the correct inducibility from the P_*petE*_ promoter was only observed in Δ*cvkR*Com-2, we employed this strain for all subsequent complementation experiments, naming it henceforth Δ*cvkR*Com.

### Deletion of alr3614 impacts *cas12k* and CRISPR array expression in vivo

The AnCAST system contains all known associated genes and extends over approximately 21 kb (**Fig. 4A**). In addition to its core CAST components, it encodes multiple other transposases, possible regulators, a toxin/antitoxin pair, and an RM system. The *cas12k* gene (*all3613*) lies upstream, reverse to *cvkR*. To analyze CvkR function *in vivo*, Northern hybridization of total RNA from triplicate clones of WT, Δ*cvkR,* and Δ*cvkR*Com with single-stranded RNA probes was performed. Specific signals for *cvkR* mRNA were detected in Δ*cvkR*Com (**Fig. 4B**). Northern hybridizations against tracrRNA and the AnCAST CRISPR array yielded signals <500 nt, which were strongly increased in Δ*cvkR* compared to the wild-type control and strongly decreased in intensity in the complementation strain Δ*cvkR*Com (**Fig. 4C**, **D**). We also detected an increased signal intensity for *cas12k* mRNA in Δ*cvkR*, while the signal was below the detection limit in WT and Δ*cvkR*Com (**Fig. 4E**), consistent with previous transcriptomic data (^19^ and Bioproject PRJNA624132). These data provide direct evidence that CvkR directly or indirectly regulates the abundance of tracrRNA promoter-derived transcript(s) and *cas12k* mRNA.

**Fig. 4.**
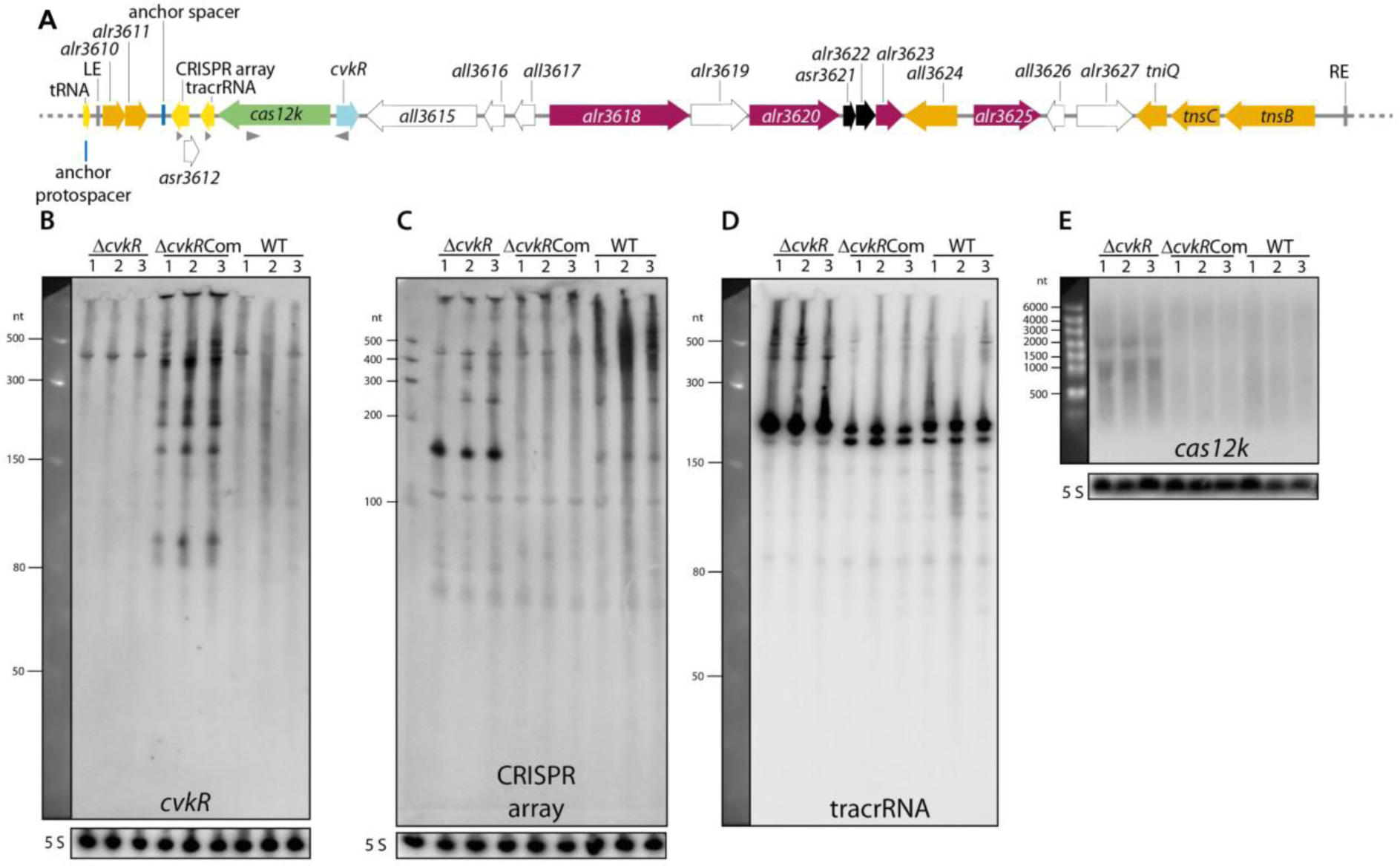
The CRISPR-associated transposase system in *Anabaena* 7120 (AnCAST) and the effects of *cvkR* deletion and overexpression on transcript accumulation. **A**. Gene arrangement within the 24,963 nt element of *Anabaena* 7120 encompassing the *cvkR* gene encoding a transcriptional regulator relative to the effector, cargo and Tn7 genes. The location of probes used for hybridizations in panels B to C is indicated by gray triangles. Gene functions are color-coded as in Fig. 1. In addition, we used pink to highlight RM genes and black to indicate a toxin-antitoxin module. **B.** Northern hybridization against *cvkR* mRNA. A signal of ∼450 nt is due to cross-hybridization, as also occurred in Δ*cvkR*. The RiboRuler Low Range RNA Ladder (Thermo Fisher Scientific) was used as a size marker. **C.** Northern hybridization against the CRISPR array (S3-S4). **D.** Northern hybridization against tracrRNA. The expected length of the tracrRNA according to transcriptome data (accession number PRJNA624132 in NCBI’s short reads archive^19^) is approximately 210 nt. The Low Range ssRNA Ladder (NEB) was used as a size marker. **E.** Northern hybridization against *cas12k*. The expected length of the *cas12k* mRNA is not known, as the transcript level was below the detection limit in WT. The gene length is 1.92 kb. RiboRuler High Range RNA Ladder (Thermo Fisher Scientific) was used as a size marker. All gels and membranes were checked for equal loading by staining with ethidium bromide and hybridization to the 5S rRNA (except in panel D because the same membrane was rehybridized as in panel B). Twenty micrograms of total RNA of the WT, the deletion mutant Δ*cvkR* and the complementation mutant Δ*cvkR*Com were separated on 10% PAA 8.3 M urea (panels B to D) or 1.5% formaldehyde-agarose gels.

Moreover, the presence of longer transcripts indicated that tracrRNA and the CRISPR array are transcribed into a longer precursor that is subsequently processed into the major accumulating fragments of ∼150 and ∼200 nt, respectively. This is consistent with previous results of differential RNA-seq^21^ and extensive transcriptome data^19^ that suggested that the AnCAST CRISPR array would not have its own specific promoter. Instead, the joint tracrRNA-CRISPR array precursor is transcribed from a TSS that is located 35 nt downstream of the *cas12k* coding sequence, at position 4,362,990 on the reverse strand, 413 nt upstream of the first repeat of the CRISPR array^14^. Nevertheless, it should be noted that the CRISPR array showed much more abundant signals in the Δ*cvkR* deletion mutant compared to WT and Δ*cvkR*Com, while tracrRNA was also well detectable in WT and Δ*cvkR*Com (**Fig. 4C**, **D**). Thus, there are additional factors involved, likely acting on the processing and stabilization of array-derived transcripts.

### Transcriptomic analysis of deletion mutant ΔcvkR and complementation strain ΔcvkRCom

To investigate transcriptomic changes upon *cvkR* deletion, microarray analysis was performed using the Δ*cvkR* and Δ*cvkR*Com deletion and complementation strains. After induction of *cvkR* transcription from the copper-inducible *petE* promoter, the mRNA levels and CvkR protein expression were verified in biological triplicate samples via Northern hybridization (**Fig. 5A**) and Western blot analysis (**Fig. 5B**). Hence, the absence or overexpression of CvkR in Δ*cvkR* or Δ*cvkR*Com, respectively, was confirmed, and the strains were investigated for further transcriptomic differences. The microarrays used cover all protein-coding genes as well as noncoding RNAs and were especially designed to allow the direct hybridization of labeled total RNA without prior conversion into cDNA^22^. Therefore, they enable the direct detection of tracrRNA and CRISPR array transcript levels. Microarray analysis revealed a small number of dysregulated genes. Thirteen features were significantly upregulated (7 protein-coding genes and 6 other transcripts) and 8 were significantly downregulated (6 protein-coding genes and 2 other transcripts) in Δ*cvkR* compared to Δ*cvkR*Com (**Fig. 5C**). In addition to the AnCAST genes *cas12k* and *tnsB* (*all3630*), the tracrRNA and CRISPR array were upregulated (**Fig. 5D**) upon deletion of *cvkR*. Hence, the role of CvkR as a regulator of the AnCAST system was confirmed and further extended.

**Fig. 5.**
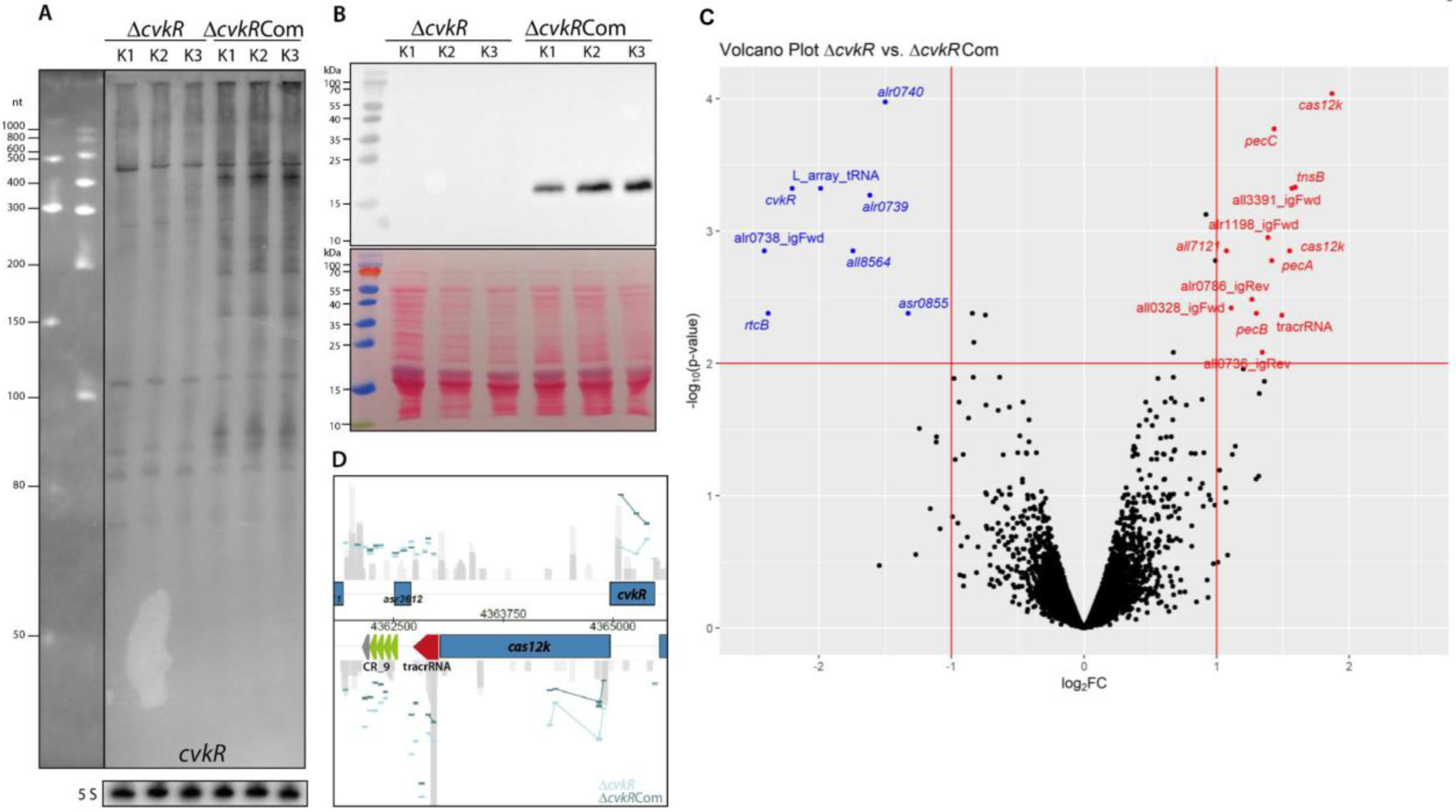
Microarray analysis of the *cvkR* deletion and complementation mutants. **A.** Northern blot hybridization against *cvkR*. The expected length of the *cvkR* mRNA is unknown because the transcript level in WT is below the detection limit. The gene length is 453 bp. Total RNA (20 µg) was loaded on a 10% PAA 8.3 M urea gel. The Low Range ssRNA Ladder (NEB) and the RiboRuler Low Range RNA Ladder (Thermo Fisher Scientific) were used as size markers. **B.** Western blot against CvkR with an N-terminal 3xFLAG tag (upper panel); the stained membrane is shown in the lower panel. The calculated molecular mass for CvkR is 20.16 kDa. The prestained PageRuler (Thermo Fisher Scientific) was used as a size marker. Ten micrograms of total protein were loaded on a 15% SDS–PAGE gel. **C.** Volcano plot of Δ*cvkR* and Δ*cvkR*Com transcriptome analyses. The horizontal line marks the p value cutoff of 0.01, while the two vertical lines mark the fold change cutoff of │log_2_│≥1. The features above these thresholds showed significant downregulation (8 transcripts, blue) or upregulation (13 transcripts, red). **D.** Visualization of the most differentially expressed region in the *Anabaena* 7120 genome, which belongs to the AnCAST system. Positions of the tracrRNA and the CRISPR array are annotated; individual probes are indicated by short horizontal bars and colored in light blue for the deletion mutant Δ*cvkR* and in dark blue for the complementation mutant Δ*cvkR*Com.

The dataset of all transcripts with meaningful fold changes is provided in **Table S2**, the raw data are available from the GEO database (https://www.ncbi.nlm.nih.gov/geo/) under the accession number GSE183629.

### CvkR DNA binding and definition of the bound sequence

To define the minimal necessary sequence and motifs required for CvkR binding, the promoter sequence of *cas12k* (P_*cas12k*_) was analyzed in a DNase I footprinting assay (**Fig. 6**). We first examined the binding activity of CvkR on a 287 bp DNA fragment covering the whole P_*cas12k*_ region in an electrophoretic mobility shift assay (EMSA). When providing equal amounts of this DNA fragment, it was with increasing concentrations of CvkR increasingly retarded (**Fig. 6A**, left panel). Then, we started to seek the CvkR binding region by further sequencing analysis. A specific area between 15 and 57 nt upstream of *cas12k* was finally identified which exhibited significant CvkR-mediated protection (**Fig. 6A**, right panel). At higher CvkR concentrations, the area expanded further even downstream of the *cas12k* start codon, which could be a result of nonspecific DNA binding or CvkR dimerization. Other areas were unaffected by CvkR addition, which indicates a specific binding affinity at this location. Interestingly, the identified DNase-protected area overlaps with an inverted repeat (IR), 5’-AAAACACA-N21-TGTGTTTT-3’ (**Fig. 6A**, right panel). To investigate this further, we also looked at other CAST systems with closely related *cvkR* genes (>70% sequence identity; **Fig. 2**) and compared their *cas12k* promoter regions, yielding six candidates. The sequences upstream of these six *cas12k* genes were aligned, and the -35 and - 10 regions of P_*cas12k*_ and P_*cvkR*_ were predicted by the PromoterHunter program^23^. The alignment of these sequences showed that the IR motifs are conserved, surround the -35 region of P_*cas12k*_ and overlap the -35 region of P_*cvkR*_ in all instances (**Fig. 6B**), which is also a typical feature of other MerR-controlled genes. Furthermore, this finding provided further support that the leaderless expression of *cas12k* and *cvkR* is conserved. Other promoters likely controlled by CvkR, such as P_*tnsB*_ and Ptracr, were also analyzed, but a closely related sequence was not found (**Fig. S2**).

**Fig. 6.**
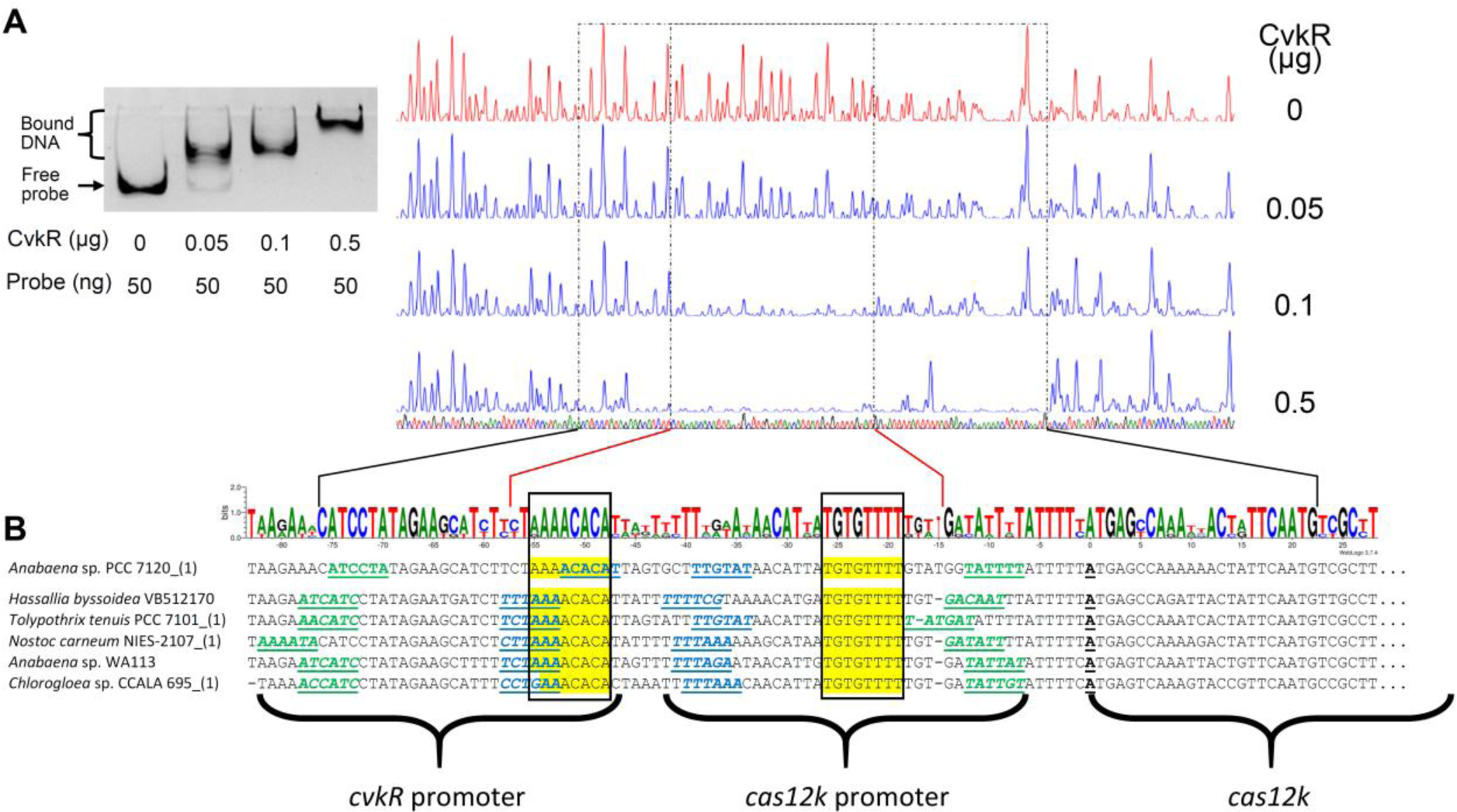
CvkR interaction with *cas12k* promoter fragments and predicted binding motifs. **A.** Left panel, EMSA showing the direct binding of different amounts of CvkR to the *cas12k* promoter DNA. Right panel, DNase I footprinting assay. The *cas12k* promoter DNA was digested in the presence of different concentrations of CvkR. The fragmentation pattern indicated a core region of 43 nt that was protected from DNase I degradation in the presence of CvkR (inner dot-and-dashed lines) embedded in a longer segment protected at higher CvkR concentrations (outer dot-and-dashed lines). **B.** The intergenic spacers between *cvkR* and *cas12k* from 6 different CASTs were aligned and analyzed for potential promoter elements. The -35 (blue) and -10 (green) regions for both promoters as well as the transcription start site of *cas12k* (black) are marked, which were previously identified^21^ or predicted by PromoterHunter^23^ (nucleotides in italics). The CvkR protected region in the DNase I footprinting assay is marked, which contains a conserved inverted repeat surrounding the -35 region of the *cas12k* promoter (boxed and highlighted in yellow). The sequences are also visualized as sequence logo.

### TXTL assays verify CvkR repressor function

To further delineate the relationship between CvkR and the promoter elements controlled by it, we tested the interaction of CvkR with different promoter fragments (**Fig. 7A**) using EMSAs and a cell-free transcription-translation system (TXTL^24^).

**Fig. 7.**
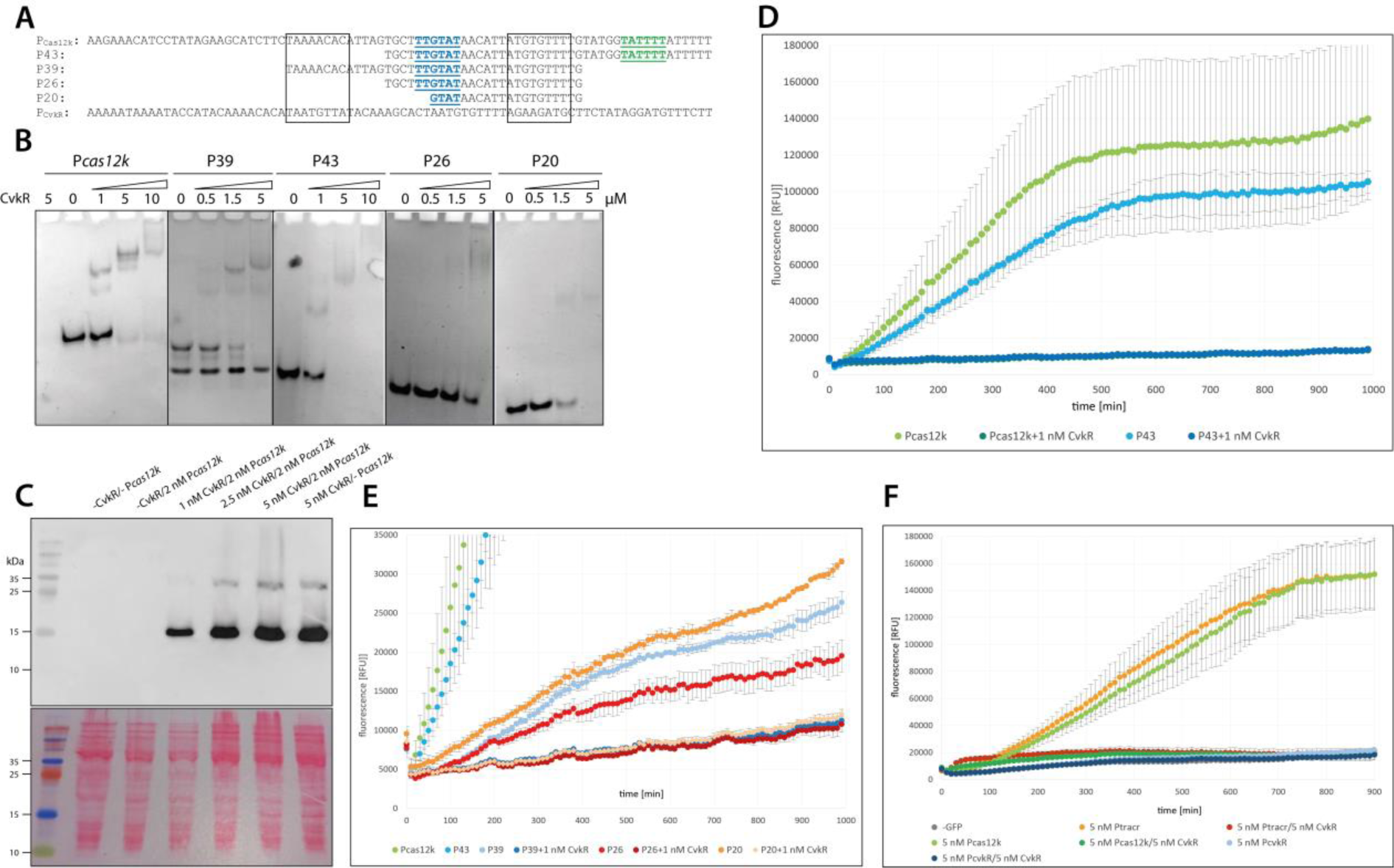
Assays to test *cas12k* promoter elements. **A.** Sequences of the tested promoter fragments. Putative -35 and -10 *cas12k* promoter elements are shown in blue and green, respectively. **B.** EMSA of the P_*cas12k*_ full-length and truncated fragments after DNA incubation for 40 min with different concentrations of CvkR. **C.** CvkR was expressed from vector pET28a and was detected by Western blot analysis via the N-terminal 6xHis tag (upper panel), and the stained membrane is shown in the lower panel. The corresponding size for 6xHis-CvkR is 19.9 kDa. The prestained PageRuler (Thermo Scientific) was used as a size marker. **D.** The full-length version of P_*cas12k*_ and the P43 fragment encompassing 43 nt upstream of the *cas12k* TSS were tested in the TXTL system^24^ for their capacity to drive deGFP expression and mediate repression upon parallel expression of CvkR. CvkR was expressed together with the corresponding p70a plasmids (5 nM) with the two promoter variants upstream of deGFP. **E.** TXTL assay for the promoter sequences P39 (positions -56 to -18 relative to the TSS of *cas12k*), P26 (positions -42 to -18) and P20 (positions -42 to -18). **F.** TXTL assay to compare the promoter activities of *cas12k*, *cvkR* and tracrRNA (P_*cas12k*_, P_*cvkR*_, Ptracr). CvkR was used to repress transcription. Error bars show standard deviations calculated from 2 technical replicates in panels D to F.

Recombinant CvkR was incubated at different concentrations with these fragments. The full-length P_*cas12k*_ promoter yielded by far the strongest interaction, and band shifts could be seen at the lowest tested concentration of 0.5 µM CvkR (**Fig. 7B**), indicating a high sensitivity for the CvkR:DNA interaction. The incubation with higher CvkR concentrations increased the signal intensity and showed a supershift or nonspecific DNA-binding affinity at the highest tested concentration of 5 µM CvkR. All other fragments showed substantially weaker interactions, independent of sequence or fragment length (**Fig. 7B**).

To test these interactions independently, we cloned the full-length *cas12k* promoter P_*cas12k*_ and several of the shorter promoter fragments upstream of a deGFP reporter gene to test whether they can drive transcription and thereby deGFP production in the TXTL assay^24^. In parallel, CvkR was expressed from a second plasmid (**Fig. 7C**).

The promoter of *cas12k* (P_*cas12k*_) was found to drive deGFP expression well (**Fig. 7D**). If CvkR was coexpressed, deGFP production was decreased to a value matching the baseline without added deGFP plasmid. P43, a 5’ shortened variant of P_*cas12k*_, yielded an ∼30% lower deGFP fluorescence than the full-length promoter and was fully repressed upon coexpression of CvkR (**Fig. 7D**). Thus, the 43 nt fragment in P43 encompassing the -10 and -35 elements and one of the palindromes constitutes a minimal promoter. Further truncation of P_*cas12k*_ on the 3’ and 5’ ends (P39, P26, P20) substantially reduced promoter activity, but the activity could still be lowered by the parallel expression of CvkR (**Fig. 7D, E**).

Next, we tested the promoter driving tracrRNA transcription (P_tracr_) in the same system. We found that P_tracr_ yielded high deGFP expression, comparable to P*_cvkR,_* and that the expression was abolished in the presence of native CvkR (**Fig. 7F**). The promoter of *cvkR* (P*_cvkR_*) was not able to drive deGFP transcription in the TXTL system (**Fig. 7F**). Therefore, whether CvkR can regulate its own transcription could not be tested in this assay.

### The crystal structure reveals that CvkR is a new type MerR-type regulator

The results shown in **Figs. 4 to 7** established CvkR as a repressor of the AnCAST system. To investigate CvkR functionality at the structural level, a high-quality crystal structure was solved at 1.5 Å resolution by using the single-wavelength anomalous dispersion (SAD) method (detailed data are in **Table S3**). A clear electron density of the added ATP molecule used in the optimization of crystallization appeared at the proposed effector-binding domain of CvkR (**Fig. 8**). However, despite our best endeavors on the apo CvkR structure, the crystal quality was insufficient for diffraction data collection. The failure might result from the high flexibility of the C-terminus without its bound effector.

**Fig. 8.**
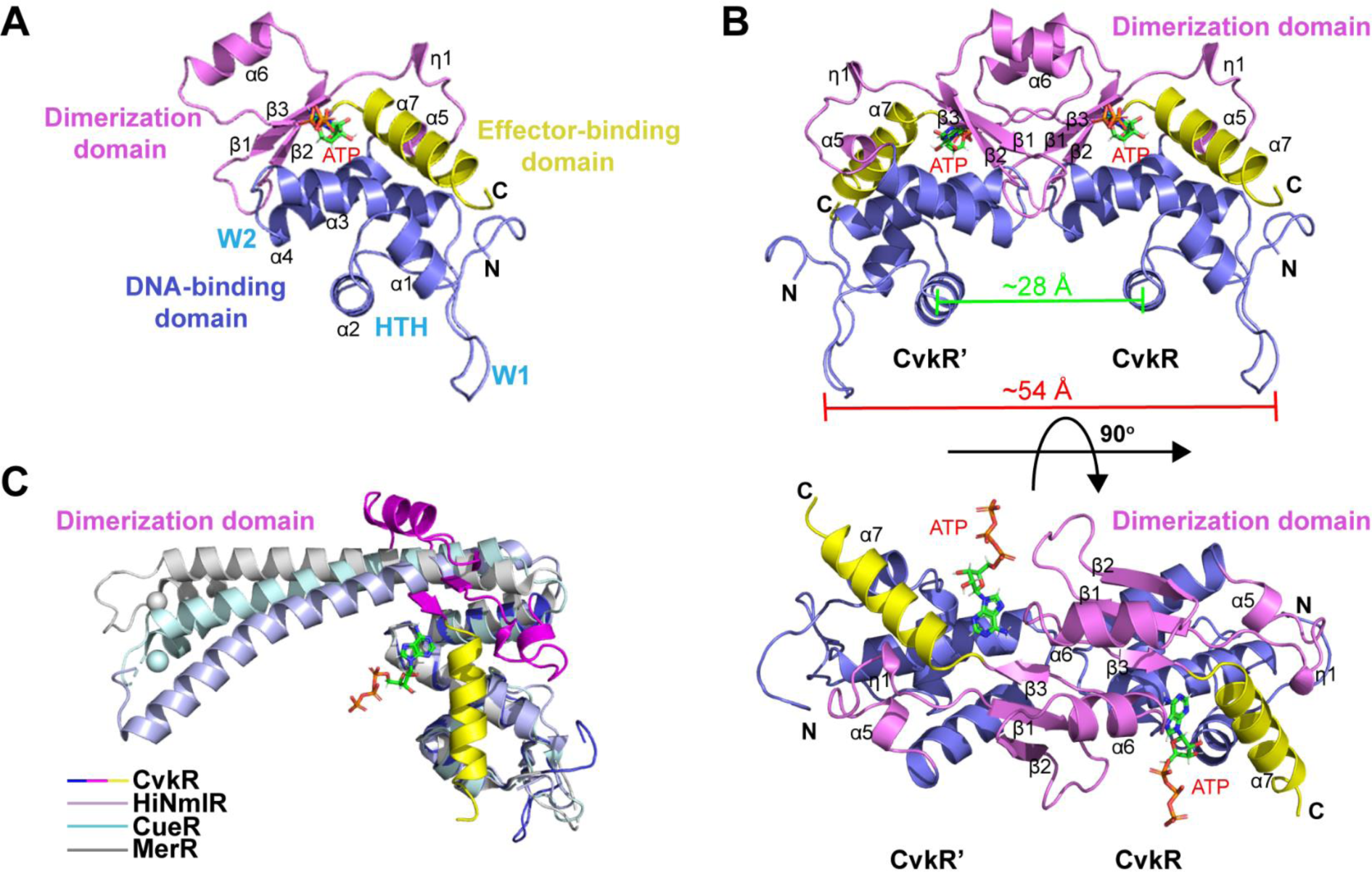
Overall structure of CvkR. **A**. Ribbon representation of the CvkR-ATP complex in ASU. The DNA-binding, dimerization, and effector-binding domains are shown in blue, magenta, and yellow, respectively. The typical helix-turn-helix (HTH) and two “wing” loops W1 and W2 in the DNA-binding domain are indicated. The ATP ligand is represented in green sticks and colored by the atom type. The secondary structure elements of CvkR are labeled. **B**. Ribbon representation of the homodimer structure of CvkR through a crystallographic symmetry operator. The novel dimerization is formed by three antiparallel β-strands (β2-β1-β3) and one short α-helix (α6) from the two protomers in the dimer. The distances between two α2-helices and between two W1 regions are measured and labeled, respectively. **C**. Structural comparison of MerR-type proteins. The DNA-binding domain of one subunit of CueR (PDB code 1Q07), MerR (PDB code 4UA1) and HiNmlR (PDB code 5D90) is separately superimposed on that of CvkR. The structural data can be accessed under the PDB accession number 7XN2.

The solved structure shows that there is one CvkR monomer binding one ATP molecule in the crystallographic asymmetric unit (ASU) (PDB ID code 7XN2, **Fig. 8A**). Further analysis through PDBePISA shows that the interface area between two adjacent CvkR monomers is as high as 970.1 Å, suggesting that CvkR forms stable homodimers in solution (**Fig. 8B**), consistent with the results of size exclusion chromatography (**Fig. S3**). Both analyses collectively indicated that CvkR functions as a homodimer similar to other reported members of the MerR family.

The overall structure of the CvkR monomer consists of a classical wHTH DNA-binding domain at the N-terminus (residues 1-80), a novel dimerization domain (residues 81-132), and a potential effector-binding helix in the C-terminal region (helix α7, residues 133-150), in which a clear ATP ligand is bound (**Fig. 8A and B**). The topology of the DNA-binding domain is typical α1-α2-W1-α3-W2-α4, which contains four α-helices and two wings and is largely preserved in the MerR-like superfamily. The classical wHTH domain structure in the CvkR N-terminus, combined with the results of our TXTL, EMSA and transcriptome analyses, confirms that CvkR belongs to the MerR-type family of transcriptional regulators. However, the dimerization domains in our solved structure differ from the existing reported MerR family members (**Fig. 8C**). Instead of dimerizing the antiparallel coiled coil formed by two longer, central α-helical linkers, CvkR dimerizes via three antiparallel β-strands (β2-β1-β3) and one short α-helix (α6) from the two protomers in the dimer (**Fig. 8B and C**). This is a completely new folding pattern of dimerization among the MerR family members reported thus far, indicating that CvkR is a structurally novel MerR-type protein.

It is well known that the C-terminal domain of MerR family regulators is responsible for specifically recognizing effector and sensing signals and ranges in size from a few residues to hundreds of amino acids. In general, the larger C-terminal domain is composed of multiple secondary structure elements and is often involved in multidrug resistance. In contrast, the shorter C-terminal domains, such as CueR and SoxR, mostly display a short α-helix structure (**Fig. 8C**) and function as bacterial sensors of metal ions or oxidative stress with the help of a conserved cysteine residue. According to the size of the C-terminal domain, CvkR belongs to the latter group; however, it lacks the conserved cysteine motif (at position 134, **Fig. S4**) and binds an ATP molecule (**Fig. 8C**). These features again indicate the novelty of CvkR in structure and function. Interestingly, the ATP ligand, used at the crystal optimization stage to obtain high-quality crystals for diffraction data collection, binds exactly to the putative effector-binding domain of CvkR (**Fig. 9**). The efficient binding of the adenine moiety is achieved through π-π stacking contributed by several aromatic residues, a cation-π interaction provided by R136, and specific hydrogen bonding patterns (**Fig. 9A**). Notably, the hydrophobic residue W133, originally embedded in the hydrophobic interior, is exposed to the solution side due to the binding of ATP, implying that some conformational changes in this region may occur with ATP binding. These interactions indicate a certain degree of base-recognition specificity, but we cannot draw a conclusion that it is adenine-specific due to the large number of water molecules participating in the formation of a hydrogen bond interaction network. The two hydroxyl groups of the ribose moiety of ATP separately formed hydrogen bonds with the side chain of residue Q140 and a water molecule. Although the interaction between Q140 and the ribose moiety also presents some specificity, the recognition specificity of this site is obviously less than that of the adenine-binding site. In contrast, the triphosphate group of ATP is free outside the CvkR molecule in our solved structure, suggesting that the ATP molecule is not the actual effector of CvkR. These findings point to an effector molecule that may be related to the cyclic oligonucleotide family of signaling molecules observed in certain types of CRISPR-Cas and other defense systems^25, 26^, but attempts testing several commercially available candidates remained inconclusive. Further structural analysis revealed that the triphosphate group of ATP was inserted into the wHTH DNA-binding domain of the adjacent CvkR’’ molecule and interacted with W1 and helix α1 (**Fig. 9B and C**). W1 in the wHTH DNA-binding domain has been proven to be involved in the binding of MerR family members to the phosphate backbone of DNA. Different from the relatively nonspecific interaction between the triphosphate group and the main chains of other residues, the side chain of residue R42 in the W1 region forms specific hydrogen bonds with the phosphate group, implying that R42 is likely to participate in the binding of CvkR to DNA (**Fig. 9C**). Therefore, the solved CvkR-ATP complex indirectly proves that CvkR binds to nucleic acids through the wHTH domain.

**Fig. 9.**
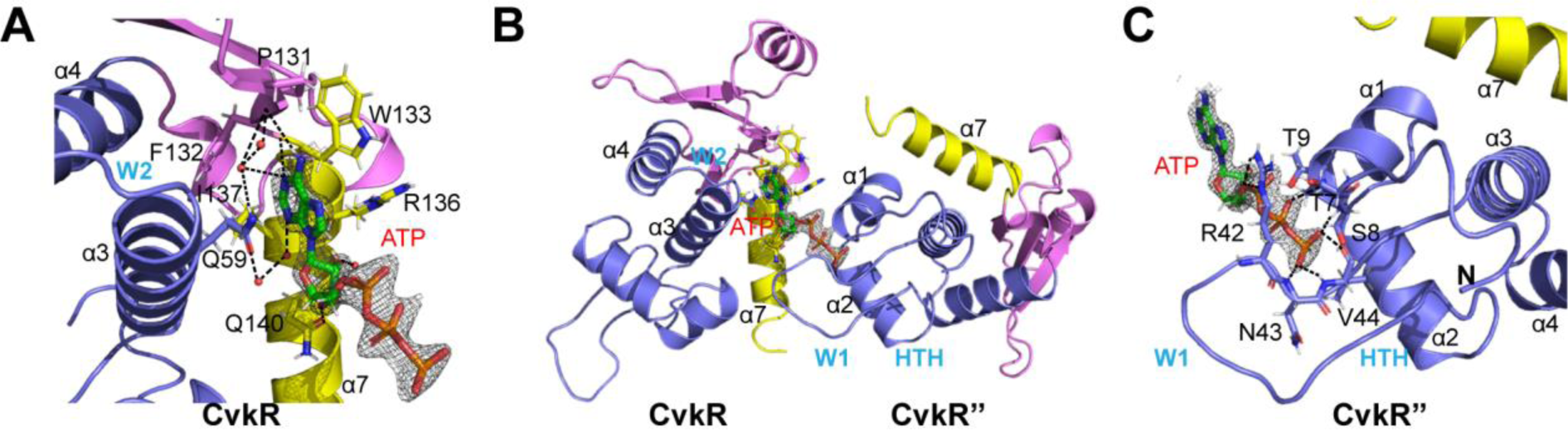
ATP binding site of CvkR. **A**. Close view of the ATP-binding site in the CvkR monomer. **B**. Overview of the ATP-binding site between two CvkR molecules (CvkR and CvkR’’). **C**. Close view of the ATP triphosphate group-binding site in CvkR’’. The ATP ligand is represented as green sticks and colored by the atom type. The 2Fo-Fc density for ATP is contoured in blue at 1.5 σ. The key residues involved in ATP binding are shown as sticks and labeled with black. The hydrogen bonds are shown by the dashed lines. The water molecules involved in hydrogen bonds are presented as red spheres.

In terms of DNA recognition, MerR regulators generally utilize residues in their α2-helix to engage the major groove, while residues in their wings of the wHTH domain engage the minor groove in the target DNA. However, the crystal quality of the CvkR-DNA complex was insufficient for diffraction data collection. Thus, we superposed our solved CvkR structure onto other reported MerR regulator-promoter complex structures. Structural comparison suggests that α2-helix residues R19, R20, Q21, Q23, Y24, R26 and E27 insert into the DNA major groove, W1 loop residues K40, R42, N43 and V44 insert into the DNA minor groove, and W2 loop residues N66, F67 and D68 are close to the DNA phosphate backbone. Except for residues R19, R26, K40, R42 and N43, the mentioned residues correspond to residues in HiNmlR, which have been found to be involved in DNA binding^27^.

To verify the relevance of these residues for transcriptional regulation, mutagenesis and TXTL functional assays were performed (**Fig. 10**). Six amino acids were changed to alanine (R19A-R20A-Q23A-K40A-R42A-N66A), and the resulting variant protein was called CvkRmut. Both CvkR and CvkRmut were stably produced in the TXTL assay (**Fig. 10A**). Similar to the assay shown in **Fig. 7**, the promoter of *cas12k* (P_*cas12k*_) was found to drive deGFP expression well (**Fig. 10B**). If CvkR was coexpressed (1 nM, 2.5 nM or 5 nM) with P_*cas12k*_-driven deGFP production, GFP fluorescence decreased to a value matching the baseline without added deGFP plasmid (**Fig. 10C**). This effect was observed at the lowest amount of added plasmid used, 1 nM. Therefore, CvkR was able to completely repress deGFP expression driven from P_*cas12k*_ at low concentrations.

**Fig. 10.**
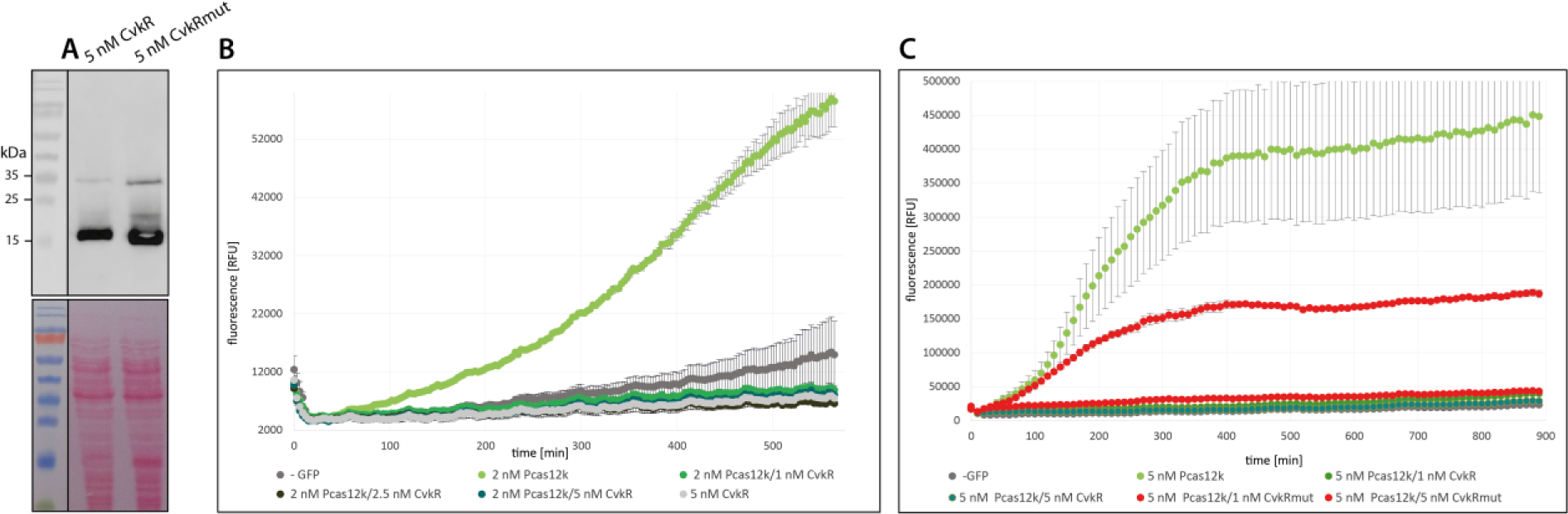
TXTL assay with CvkR and CvkRmut and different promoters. **A.** To address the relevance of possibly critical residues according the CvkR crystal structure (Fig. 8 and 9), six amino acids were substituted by Ala (R19A-R20A-Q23A-K40A-R42A-N66A), yielding protein CvkRmut. *Anabaena* 7120 CvkR and CvkRmut were expressed in the TXTL system^24^ from pET28a and detected by Western blot analysis via their N-terminal 6xHis tag (upper panel). The stained membrane is shown below. The corresponding molecular masses are 19.9 kDa for 6xHis-CvkR and 19.5 kDa for 6xHis-CvkRmut. **B.** The promoter of *cas12k* (P_*cas12k*_) was used to express deGFP from plasmid p70a (2 nM) in the absence or presence of different amounts of plasmid pET28a expressing CvkR. Error bars show the standard deviation and are derived from two technical replicates. **C**. TXTL assay to test the regulatory capacity of CvkRmut compared to CvkR for repressing deGFP fluorescence expressed from p70a under the control of P_*cas12k*_ (5 nM). Error bars show the standard deviation and are derived from 2 technical replicates. All experiments were repeated twice independently. The TXTL reactions were performed at 29 °C overnight, and fluorescence was measured every 10 min in a Wallac 1420 Victor 2 microplate reader.

In contrast, CvkRmut expressed from the 1 nM plasmid could not repress P_*cas12k*_-driven deGFP expression as efficiently as CvkR. However, if a higher amount of 5 nM plasmid was added, CvkRmut was able to repress deGFP almost to the same level as CvkR expressed from 1 nM plasmid (**Fig. 10C**). These results indicated that CvkRmut was a mild binding-deficient mutant that was still functional but not as efficient as the native CvkR, hence supporting the predicted functional relevance of these residues.

## Discussion

Native type V-K CAST systems have thus far only been found in certain cyanobacteria^11, 14^. Their better characterization is of fundamental interest, and these systems hold great promise for the development of novel genome editing tools ^28^. The primary function of native CRISPR-Cas systems is defense against mobile genetic elements. Therefore, we reasoned that the activity of CAST systems that can transpose to novel sites within a genome must be controlled. Here, we first addressed the association between putative transcriptional regulators and cyanobacterial CAST systems and found four different classes of such repressors, which we suggest to name CvkR, for Cas-type V-K repressors. The consistent occurrence of these regulators in 94 of 118 analyzed systems indicates a functional dependency. Judged by the diversity of regulators and their DNA-interference domains, CvkRs likely have originated several times independently.

We then characterized the transcriptional regulator of the AnCAST system of *Anabaena* 7120 encoded by gene *alr3614* in detail. Our data show that CvkR controls the expression of the AnCAST core module (i.e., tracrRNA, *cas12k* and *tnsB*, *tnsC* and *tniQ* mRNAs).

The crystal structure analysis revealed specific features within the dimerization and effector-binding domains of CvkR that were previously unknown for members of the MerR-type family of transcription factors. The MerR family was first discovered as a regulator of mercury resistance operons^29, 30^ but is also involved in multiple cell functions, such as drug resistance, responses to heavy metals and protection against oxidative stress^31–33^. In gram-negative bacteria, transposable elements such as Tn21 and Tn501 frequently contain mercury resistance (*mer*) operons, which are controlled by transcription factors belonging to the MerR family.

MerR regulators typically contain an N-terminal HTH domain followed by a dimerization helix and an effector-binding domain at the C-terminus^29, 34^. The protein forms a coiled-coil homodimer facilitated over the dimerization helix and is able to bind a palindromic DNA motif^35^. The protein can bind DNA with and without effector binding, which leads to the protein functioning as either an activator or a repressor. The palindromic binding region is usually inside promoters with an elongated distance (>19 nt) between the -10 and -35 elements, so MerR represses the binding of the σ-factor^29^. This is also true for the AnCAST system and CvkR, in which the distances between the -35 and -10 elements are 21 nt for the *cas12k* promoter (**Fig. 6B**) and 19 nt for the Ptracr promoter. In contrast, this distance is 17 nt for the *cvkR* and *tnsB* (*all3630*) promoters.

Effector binding, however, can trigger a conformational change, which brings both HTH domains closer together and distorts the DNA to increase the σ-factor affinity for the promoter^29, 35^. The effector for this regulator is usually a metal ion, but there are also MerR interactions with antibiotics, oxidative stress, or lipophilic compounds^29, 36–38^. MerR-like proteins without effector interactions are also known^39^.

No fluorescence was observed if CvkR was coexpressed in a cell-free transcription-translation system (TXTL) together with deGFP reporter gene fusions of the *cas12k* promoter P_*cas12k*_ and the tracrRNA promoter Ptracr, while in the absence of CvkR, strong signals were detected (**Fig. 7**). These results establish CvkR as a transcriptional repressor of the AnCAST system. We tested mutated versions of the protein and of the promoter sequence, yielding insight into the functionality of the protein as a dimeric HTH-domain-containing transcriptional repressor and allowing delineation of the binding motif at the DNA level. The genome-wide analysis of transcriptomic effects yielded, in addition to the three CvkR-controlled promoters shown in **Fig. 7F**, evidence for further promoters possibly under its control, both on the chromosome and on plasmids α and δ (**Fig. 5C** and **Table S2**). MerR-type regulators are principally better known as transcriptional activators rather than transcriptional repressors. Our data show that transcription of three promoters in the AnCAST system was enhanced in the Δ*cvkR* mutant. While we detected the repressor function as the default mechanism, we speculate that this interaction might become lost in the presence of a bound effector molecule.

Genes upregulated in Δ*cvkR*Com (**Fig. 5C**) include a set of 26 tRNA genes on the ∂ plasmid. These were recently identified as an L-array that became specifically induced when cultures were exposed to sublethal concentrations of ribosome-targeting antibiotics^40^. Moreover, the adjacent gene *all8564* encoding an HNH-type homing endonuclease and the chromosomal gene *all3626* (*rtcB*) were found to be coregulated. Therefore, the genes that were significantly more highly expressed in Δ*cvkR*Com indicated the presence of translational stress due to the presence of antibiotics.

Structural analysis of CvkR at 1.5 Å resolution yielded elements typical for a member of the MerR-type transcription factor family but with distinct dimerization and effector-binding domains. In particular, the 28 Å and 54 Å distances between two α2-helices and two W1 loops in CvkR, covering ∼9 bp and ∼17 bp, respectively, in B-form DNA, are both obviously shorter than those in the HiNmlR-promoter complex (35 Å/74 Å;^27^) and other reported MerR regulator-promoter complexes, such as SoxR (35 Å/74 Å) and the activator CopA (35 Å/65 Å) (**Fig. 8B**). The α2-helix in the wHTH DNA-binding domain is well known for base-specific recognition by MerR-type transcriptional regulators. Individual MerR-family homodimer proteins bind specifically to the two half-sites of quasi-palindromic inverted repeat (IR) DNA sequences within the target gene promoter via its two α2-helices. This is compatible with the length of one IR half-site recognized by CvkR of 8 bp (**Fig. 6B**).

To summarize, we identified and characterized CvkR as the regulator of the AnCAST system and found that it may impact a small set of host genes. Structural analysis of CvkR revealed that it is a structurally novel type of MerR protein because it dimerizes via three antiparallel β-strands and one short α-helix (α6) (**Fig. 8B and C**). CvkR exhibits an effector-binding domain in its C-terminal region (**Fig. 9**), and the efficient binding of an adenine moiety points to a metabolite that may be related to the cyclic oligonucleotide family of signaling molecules observed in certain types of CRISPR-Cas systems^25^, as well as other antiphage signaling systems^26^. However, the exact effector molecule and the functionality of this hypothetical signaling input sensed by CvkR are matters of further research.

## Materials and methods

### Cultures of cyanobacteria and construction of mutant strains

*Anabaena* 7120 and its derivatives were grown photoautotrophically in BG11 liquid medium or on agar plates under white light illumination of 30-50 µmol photons m^-2^ s^-1^ at 30 °C^41^. In terms of the strains for the control group in copper-inducible experiments, transparent plastic tissue culture flasks were used for cultivation, deionized water was used for medium preparation, and three resuspensions with copper-free medium were employed for the seed culture. In addition, excess CuSO4 (1.0 or 1.25 μM) was added to guarantee P_*petE*_ activity for the respective experiments. The culture was supplemented with erythromycin (10 μg/mL) when necessary.

For construction of *Δalr3614* (Δ*cvkR*), the CRISPR-Cas12a (Cpf1) genome editing tool together with the pSL2680 plasmid (Addgene No. 85581) were used as previously described^19^. The primer pair alr3614gRNA-1/2 was used to prepare the gRNA-cassette editing plasmids, and the primer pairs alr3614KO-1/2 and alr3614KO-3/4 were used to prepare the gRNA & repairing-cassette editing plasmids. The primer pairs alr3614-3/4 were used to check the deletion genotype. A 200 bp internal fragment (4365123∼4365322) of *cvkR* was ultimately deleted.

For complementation of the Δ*cvkR* mutant and/or verification of leaderless expression of the *cvkR* gene, three cassettes (P_*petE*__no 5’UTR-*alr3614*L-3xFLAG, P_*petE*__no 5’UTR-*alr3614*S-3xFLAG and P_*petE*_-*alr3614*S-3xFLAG) were cloned into a shuttle vector (pRL59EH) derived from the broad-host-range plasmid RSF1010 by seamless assembly. The primer pairs 59M-F/59M-R1, P_*petE*_-F/P_*petE*_-R4 and 3614L-F/3614-R were used to construct a complementation plasmid to generate Δ*cvkR*Com-1. The primer pairs 59M-F/59M-R1, P_*petE*_-F/P_*petE*_-R5 and 3614S-F1/3614-R were used to construct a complementing plasmid to generate Δ*cvkR*Com-2. The primer pairs 59M-F/59M-R1, P_*petE*_-F/P_*petE*_-R6 and 3614S-F2/3614-R were used to construct a complementing plasmid to generate Δ*cvkR*Com-3. Each complementing plasmid was introduced into the Δ*cvkR* mutant by conjugal transfer as previously reported^42^. Genotypes of mutants were confirmed by PCR (**Fig. S1**). The sequences of all oligonucleotides are listed in **Table S4**. All PCR fragments, plasmids generated in this study, and gene mutation regions in the mutants were verified by Sanger sequencing.

### Microarray analysis

*Anabaena* 7120 strains Δ*cvkR* and Δ*cvkR*Com were grown in 50 mL BG11 without CuSO4 to an OD750 of 0.8, and CvkR expression was induced from P_*petE*_ with 1.25 µM CuSO4 for 24 h. Cells were harvested, and RNA was extracted as described^19^. The RNA samples of two biological replicates each were hybridized to 8x44K microarrays (Agilent ID 062842) following published sample preparation and hybridization details^43^. In short, 2 µg of DNase-treated RNA was used for Cy3 labeling (ULS Fluorescent Labeling Kit for Agilent Arrays, Kreatech). Microarray hybridization was performed with 600 ng Cy3-labeled RNA for 17 h at 65 °C. Microarray raw data were processed with R software as described^22^. A |log_2_ FC|≥ 1 threshold and a p value ≤ 0.01 were considered to indicate a significant change in gene expression. The full dataset is accessible in the GEO database (https://www.ncbi.nlm.nih.gov/geo/) with the accession number GSE183629.

### Northern blot analysis of mutants

RNA isolation was performed using a Precellys 24 Dual homogenizer (Bertin) for cell lysis as previously described^19^. Twenty micrograms of total RNA were separated on 10% polyacrylamide-8.3 M urea gels. CRISPR-related transcript accumulation was analyzed by Northern hybridization using single-stranded radioactively labeled RNA probes transcribed *in vitro* from PCR-generated templates (see **Table S4** for primers), as previously described^44^.

### Western blot analysis

Cyanobacterial cell harvesting and protein extraction were performed as previously described^45^. Total proteins extracted from the samples were separated by 15% SDS– PAGE according to the standard procedure and electroblotted onto PVDF or nitrocellulose (NC) membranes. To check for equal loading, the membrane was stained with Ponceau S (0.1% (w/v) in 5% acetic acid). After destaining, the PVDF membranes were blocked in 3% skimmed milk-TBST (0.05% Tween-20 in TBS) at room temperature for 30 min. Then, the membranes were incubated with anti-3xFLAG tag monoclonal antibodies for 1 h and washed three times with TBST (15 min each). After that, the membranes were incubated with an alkaline phosphatase-linked secondary antibody for 1 h and washed three times with TBST (15 min each). Finally, signals were detected with the NBT (nitro-blue tetrazolium chloride) and BCIP (5-bromo-4-chloro-3’-indolyphosphate p-toluidine salt) methods. Methods for Western blot analysis of TXTL samples were used as described^19^.

### Quantitative real-time PCR

Total RNA extraction, removal of the genomic DNA and reverse transcription were performed using a Bacteria RNA Extraction Kit and HiScript III RT Supermix for qPCR (+gDNA wiper) kit (Vazyme) according to the manufacturer’s instructions. SYBR Premix ExTaqTM (Takara, Dalian, China) was used for qRT–PCR, and the cycle thresholds were determined using a Roche LightCycler® 480 II sequence detection system (Roche, Shanghai, China). *rnpB* (RNase P subunit B) was used as the internal control. The primers for *alr3614* (*cvkR*) and *rnpB* are listed in **Table S4**. Three independent experiments were performed, which showed consistent results.

### TXTL assays

To test promoter fragments in a cell-free transcription-translation system, the *E. coli*-based TXTL assay was used^24, 46^. The myTXTL Sigma 70 Cell-Free Master Mix was purchased from Arbor Biosciences. The included p70a plasmid was used as a template for cloning of the promoter sequences Ptracr, P_*cvkR*_, and P_*cas12k*_ (**Fig. 7B**) in an open reading frame with the destabilized enhanced GFP (deGFP) and its 5’UTR. All PCRs were performed using PCRBio HiFi polymerase (PCR Biosystems). Promoter sequences were PCR-amplified from genomic DNA of *Anabaena* 7120 with overlaps to p70a. The p70a plasmid was also PCR-amplified.

The CvkR protein (Alr3614) and a mutant with potential reduced DNA-binding affinity (R16A-R20A-Q23A-K40A-R42A-N66A) (CvkRmut) were used for analysis in the TXTL assay. The *cvkR* sequence was PCR-amplified from genomic DNA of *Anabaena* 7120. The corresponding DNA fragment for CvkRmut was ordered from IDT as gBlocks and subcloned into pJet1.2/blunt (Thermo Fisher Scientific) and then amplified via PCR with overhangs to pET28a. The plasmid pET28a was PCR-amplified as well. The CvkR and CvkRmut proteins were thus expressed from an IPTG-inducible T7 promoter with an N-terminal 6xHis tag and a TEV site for potential cleavage of the tag.

Fragment assembly via AQUA cloning was performed at room temperature for 30 min upon transformation into chemically competent *E. coli* DH5α cells for cloning. Assembled plasmids were isolated, and regions of interest were sequenced (Eurofins Genomics).

For the expression of proteins encoded on pET28a in the TXTL assay, T7 RNA polymerase (RNAP), expressed in this instance from p70a, and IPTG are necessary. Reactions were performed in duplicate overnight at 29 °C in a total volume of 5 µL with 3.75 µL of TXTL master mix, 1 mM IPTG, 0.5 nM p70a_T7_RNAP, 1 to 5 nM pET28a_CvkR or CvkRmut and 2-5 nM p70a_promoter_deGFP. The fluorescence of deGFP was measured every 10 min (excitation 485 nm, emission filter 535 nm) in a plate reader (Wallac 1420 Victor^2^ microplate reader from Perkin Elmer). Western blot analysis of TXTL samples was performed as described^19^.

### Heterologous expression and purification of CvkR protein

For heterologous expression of CvkR in *E. coli*, the protein-coding sequence of CvkR was PCR-amplified from *Anabaena* 7120 genomic DNA and cloned into the pET28a-smt3 vector by using BamHI and XhoI restriction sites to generate the pET28a-smt3_CvkR expression plasmid, which expresses CvkR with an Ulp1-cleavable N-terminal 6xHis-smt3 fusion tag. The used primers are listed in **Table S4**. The sequence-verified plasmid was then transformed into *E. coli* BL21(DE3) for protein expression.

The expression strain was grown in LB medium to approximately OD600∼0.6 at 37 °C and induced at 25 °C overnight with 0.4 mM IPTG (isopropyl β-D-thiogalactopyranoside). After induction, cells were harvested by centrifugation and lysed in lysis buffer A (20 mM Tris-HCl, pH 8.0, 300 mM NaCl, 10 mM imidazole, 2 mM β-mercaptoethanol, 20 μg/mL DNase I) using a high-pressure homogenizer. After centrifugation at 20,000x g for 50 min, the supernatants were loaded onto a Ni Sepharose 6 FF column (GE Healthcare), washed with lysis buffer A containing 40 mM imidazole, and eluted with 20 mM Tris-HCl, pH 8.8, 300 mM imidazole, and 2 mM β-mercaptoethanol. The eluted fractions were treated with Ulp1 protease overnight at 4 °C and purified with a Hitrap Heparin HP column (GE Healthcare) to remove the 6xHis-smt3 fusion tag and impurities. Subsequently, tag-removed target protein fractions were further purified by Superdex 200 Increase 10/300 GL (GE Healthcare). Finally, target proteins were collected, concentrated and stored in 10 mM Tris-HCl (pH 8.8), 100 mM NaCl, and 2 mM β-mercaptoethanol.

Selenomethionine-labeled (Se-Met) CvkR protein was overexpressed and purified using the same procedures described above; however, the medium was substituted with 1x M9 medium, and seven essential amino acids were added at the mid-log phase before induction. To prevent oxidation of the selenium atoms, 5 mM β-mercaptoethanol was added to the final elution fraction containing Se-Met CvkR.

### Analysis of CvkR oligomeric forms in solution

Size exclusion chromatography was performed with 0.3 mg CvkR at room temperature to probe the molecular weight of CvkR in solution. The Superdex 200 increase 10/300 GL column was calibrated with a gel filtration calibration kit HMW (GE Healthcare) in a buffer containing 10 mM Tris-HCl (pH 8.8), 100 mM NaCl, 2 mM β-mercaptoethanol. The calibration curve based on the molecular markers is logMr=-0.20945Ve+7.712 (R^2^=0.9949, Ve: elution volume; Mr: molecular weight).

### EMSA analysis

Each DNA duplex for EMSA was created by annealing two complementary oligonucleotides. The EMSA reaction (10 μL) was carried out at room temperature by mixing 0.5 μM DNA duplex and increasing concentrations of CvkR protein in binding buffer (50 mM Tris-HCl, pH 8.0, 100 mM KCl, 2.5 mM MgCl2, 0.2 mM DTT, 10% glycerol). After incubation for 30 min, the reaction samples were electrophoresed on an 8% polyacrylamide gel with 0.5x TBE, and the gel was visualized by ethidium bromide staining.

### DNase I footprinting assays

DNase I footprinting assays were carried out similar to previous research^47^. Specifically, ∼300 nt FAM-labeled probes, including an 82 nt intergenic spacer between the *cas12k* and *cvkR* genes, were PCR amplified with 2x TOLO HIFI DNA polymerase premix (TOLO Biotech, Shanghai) using primers P3613-F(FAM) and P3613-R and purified by the Wizard® SV Gel and PCR Clean-Up System (Promega, USA). Binding reactions were performed in a total volume of 40 µL containing 50 mM Tris-HCl, pH 8.0, 100 mM KCl, 2.5 mM MgCl2, 0.2 mM DTT, 10% glycerol, 2 μg salmon sperm DNA, 300 ng probes and 0.1 μg CvkR protein at room temperature for 30 min. Following DNase I treatment (Promega), phenol/chloroform extraction, and ethanol precipitation, products were dissolved in 30 µL MiniQ water. The preparation of the DNA ladder, electrophoresis and data analysis were the same as described before^47^, except that the GeneScan-LIZ600 size standard (Applied Biosystems) was used.

### Phylogenetic analysis of CvkR

The identified CvkR proteins were compared to each other and analyzed for alternative start positions to correct potential incorrect annotations, as in the case of CvkR (Alr3614, BAB75312.1). Elongated N-terminal regions with no similarity to homologs encoded by other *cvkR* genes were removed from the analysis. The sequences were then aligned by M-coffee^48, 49^ and further analyzed by the BEAST algorithm^50^. The phylogenetic analyses were calculated by the Yule process (speciation), the substitution model Blosum62 and MCMC chain length of 1e6 to log parameters every 1e3 steps^51–53^.

### Crystallization, data collection, and structure determination of the CvkR protein

Using a hanging drop-vapor diffusion method, both native and selenomethionine-substituted (Se-Met) CvkR crystals appeared at 18 °C in crystallization reagent containing 0.1 M phosphate citrate (pH 4.4∼4.6) with 15%∼20% PEG300. However, diffraction-quality crystals were obtained only when 10 mM ATP was added into the crystal screen droplet, which consisted of a 1:1 (v/v) protein at 15 mg/mL and the well crystallization reagent. Before flash cooling in liquid nitrogen, crystals were cryo-preserved using crystallization reagent supplemented with 15% PEG400.

The diffraction data were collected at the Shanghai Synchrotron Radiation Facility (SSRF), beamlines BL17U1 and BL18U1, in a 100K nitrogen stream. Data indexing, integration, and scaling for native CvkR were conducted using HKL3000 software^54^. SAD X-ray diffraction data were processed by Aquarium software^55^. The resulting main chain structure was used as the initial search model for molecular replacement by Phaser to determine the native CvkR structure^56^. Structure refinements were iteratively performed using the programs Phenix and Coot^57, 58^. The statistics for data processing and structure refinement are shown in **Table S3**. The coordinates were deposited in the Protein Data Bank with the PDB ID code 7XN2. Figures were prepared using PyMOL^59^.

## Data availability

The full transcriptome datasets for the WT and mutants Δ*cvkR* and Δ*cvkR*Com are accessible from the GEO database (https://www.ncbi.nlm.nih.gov/geo/) with the accession number <u>GSE183629</u>. The structural data can be accessed at the Protein Data Bank (https://www.rcsb.org/) under the PDB accession number 7XN2.

## Author contributions

WRH, XL and TZ designed the study. TZ, YX and HL constructed the *Anabaena* 7120 mutant strains. VR performed the TXTL experiments, microarray analyses and Northern hybridizations. MZ did the majority of bioinformatic analyses. YS and YX performed the qRT–PCR and Western blot analyses. YL performed the structural analysis of CvkR. VR, MZ, YL, TZ and WRH wrote the paper with contributions from all authors.

