## Supplemental Figures S1 to S4 and Tables S2 to S4 for "CvkR, a novel MerR-type transcriptional regulator, is a repressor of class 2 type V-K CRISPR-associated transposase systems"

**Supplemental Material**

### Supplemental Figures

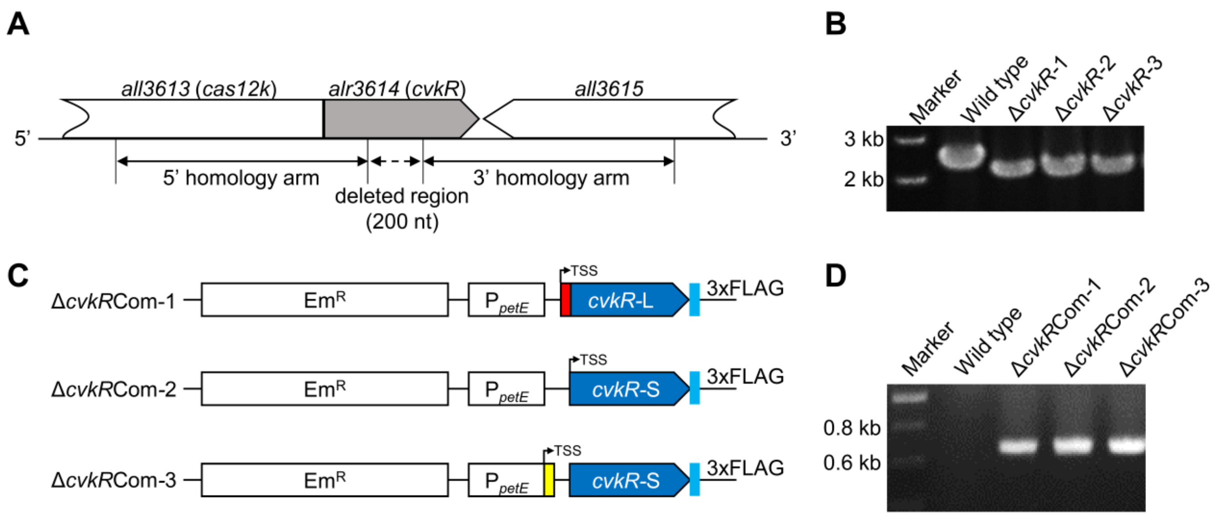


**Figure S1. Generation of *cvkR* deletion mutants and complementation of *ΔcvkR*.** **A.** Schematic drawing of the deleted regions and flanking regions of *cvkR*. **B.** PCR assays showing successful segregation of deletion mutants in three independent replicates each. **C.** Schematic drawing of the three cassettes for analyzing the leaderless expression of *cvkR*. **D.** PCR assays showing successful construction of reporter strains for monitoring leaderless expression of *cvkR*.

## A

#### Upstream *tnsB*/*tnsB*-Pseudogenes(*):

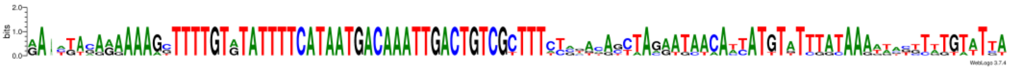

```
Anabaena sp. PCC 7120_(1)*        AA-TTACAAAAAAGGTTTTGTATATTTTCATAATGACAAATTGACTGTCGTTTTCTCCACGTTTAGAATAACATTATGTATTTATAAATTACCTTTGTTTCA

Hassallia byssoidea VB512170      AA-ATAGAAAAAAGCTTTTGTGTATTTTCATAATGACAAATTGACTGTCGCTTTATCTACACCTAGAATAACAATATGTATTTATAAAAAACTTATGTATTA
Tolypothrix tenuis PCC 7101_(1)*  GAACGACAGAAAAGCTTTTGTGTATTTTCATAATGACAAATTGACTGTCGCTTTCTGGGGAGCAACGTGCTAACCATTTATCGGCAAGTGGGTTGGGTATTT
Nostoc carneum NIES-2107_(1)      AA-CGACAGAAAAGCTTTTGTGTATTTTCATAATGACAAATTGACTGTCGCTTTCTGTATACCTAGAATAACAATATGTATTTATAAAAAAGTTATGTATTA
Anabaena sp. WA113                GA-TT-AAAAAAAGTTTTGTATATTTTCATAATGACAAATTGACTGTCGCTTTCCAGTAAGCTAGAATAACATCATGTTTTTATAAAATCTGTTTGATTTA
Chlorogloea sp. CCALA 695_(1)*    GA-ATTGGGGAAAAGTTTTGTATATTTTCATAATGACAAATTGACTGTCGCTTTTCAAACAGTTAAGATAACATTATGTGTTTATAAAGTTTGCTTTTATGA
```

*tnsB* promoter

## B

#### Upstream tracrRNA:

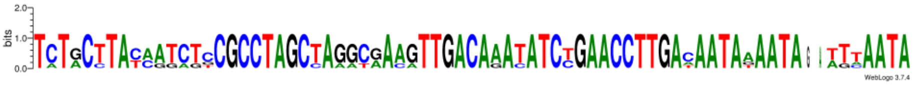

```
Anabaena sp. PCC 7120_(1)         TATGCTTATAGTCTCCGCCTAGCTAGGCGAAGTTGACAAACATCTGAACCTTGAAAATATAATA--TGTAATA

Hassallia byssoidea VB512170      TCTGCTTACAATCTTCGCCTAGCTAGGCGAAGTTGACAAATATCTGAACCTTGACAATAAAATA--ATAAATA
Tolypothrix tenuis PCC 7101_(1)   TCTACCTACCATAGCCGCCTAGCTAGGTGACATTGACAAATATCTGAACCTTGACAATAAAATA--ATCAATA
Anabaena sp. WA113                TCTACTTATCATCTCCGCCTAGCCAAACAAAGTTGACAAATATCCGAACCTTGACAATAAAATAGATTTAATA
Chlorogloea sp. CCALA 695_(1)     TCTGCTTACAAGCTGCGCCTAGCTAGGCAAAGTTGACAGATATCCGAACCTTGATAATAGAATA--TTTAATA
```

tracrRNA promoter

**Figure S2. Multiple sequence alignment of the promoters of *tnsB* and tracrRNA.** The promoter areas of *tnsB* (**A**) and tracrRNA (**B**) from cyanobacterial CASTs with closely related *cvkR* genes were aligned and analyzed for their potential -35 (blue) and -10 (green) regions (previously identified, or in italics, predicted by PromoterHunter[1]). *Nostoc carneum* NIES-2107 lacks a tracrRNA gene, so only 5 trancrRNA sequences were compared.

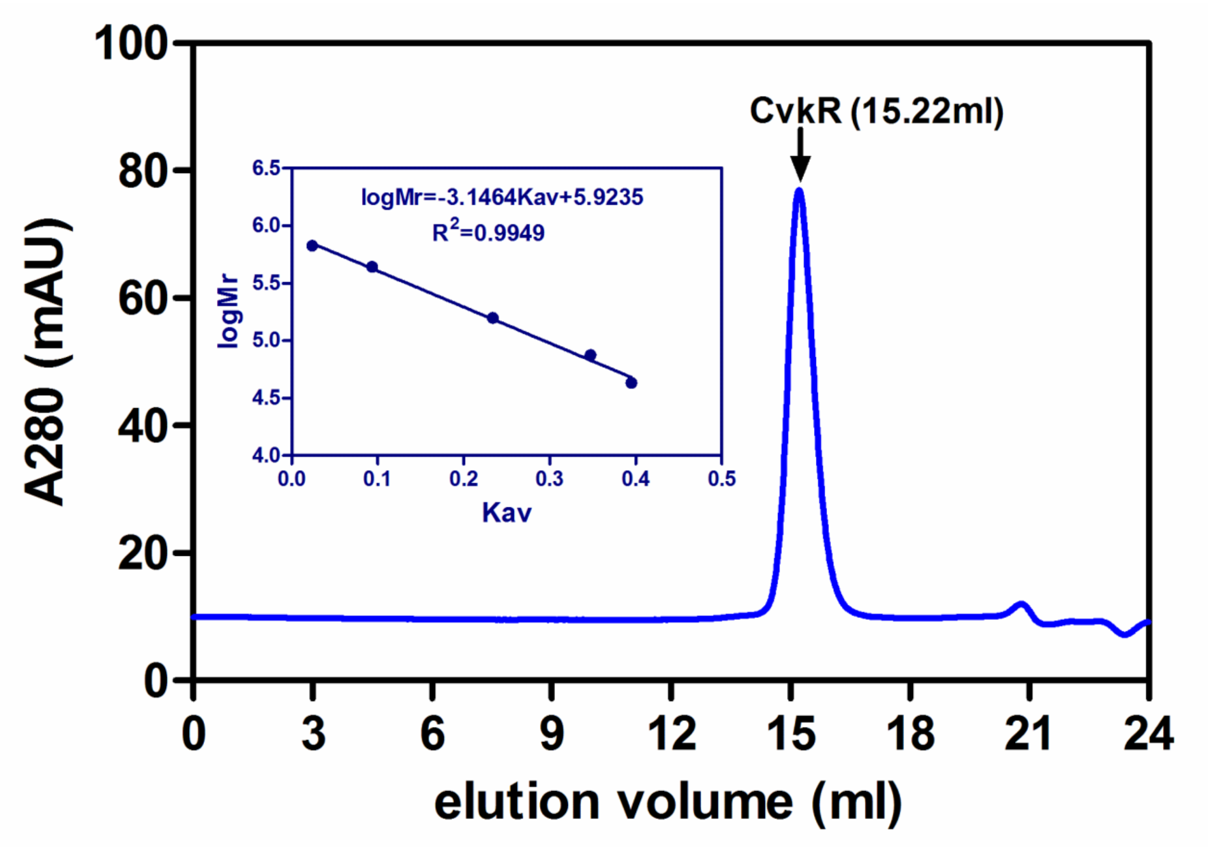


**Figure S3. Molecular weight estimates of CvkR on SEC.** X axis, elution volume (ml); Y axis, absorption at 280 nm (mAU, milliabsorbance units). The calibration curve (plot of logMr versus Kav) based on molecular markers is shown in inset. Gel phase distribution coefficient Kav=(Ve-8.54)/15.022, Ve: elution volume.

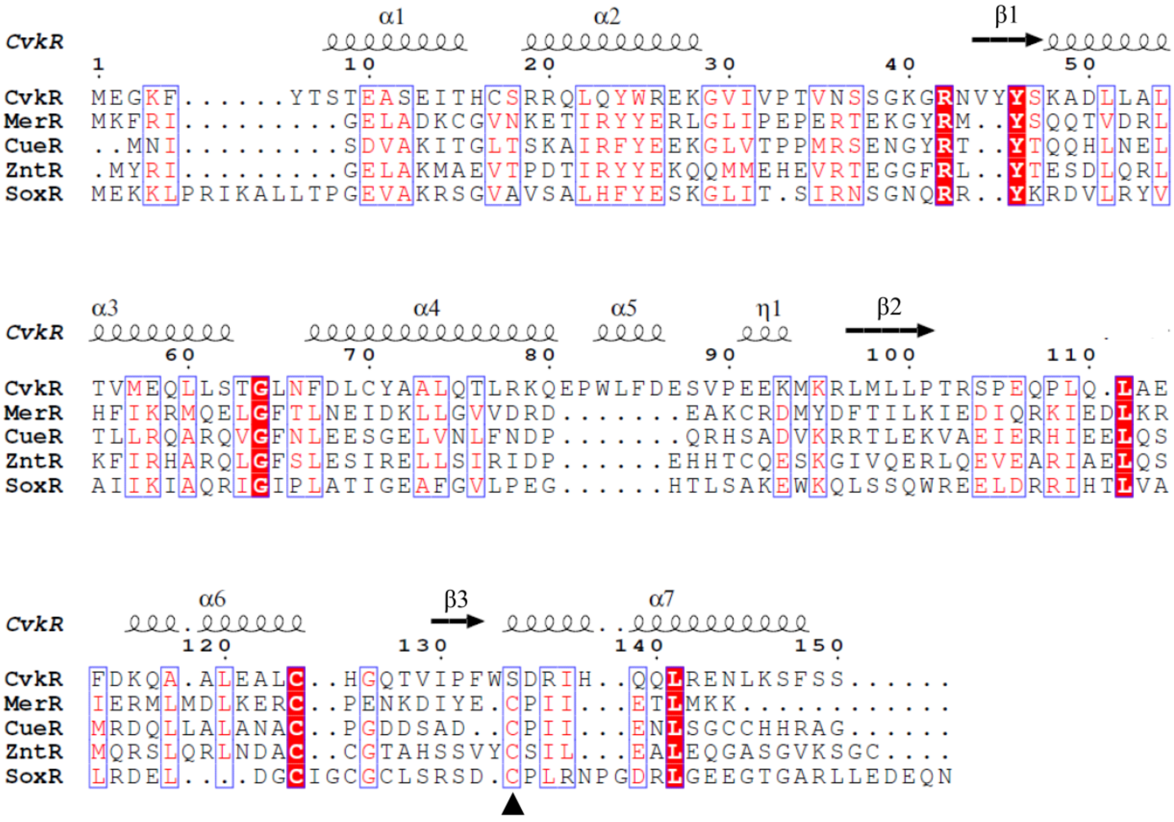

**Figure S4**. **Multiple sequence alignment of CvkR with selected members of the MerR-type regulators.** The secondary structure information of CvkR was labeled with scribble (α-helices) and black arrows (β-sheets), respectively. CvkR comes from *Anabaena* 7120; MerR is metal-dependent MerR-like regulators from *Bacillus megaterium.* CueR and ZntR are metal-dependent MerR-like regulators from *E. coli*. SoxR is oxidative stress response regulator from *E. coli*. Sequence identities are highlighted in red and similarities are displayed as red letters. Sequence analysis was performed using MAFFT[2], and the figure was prepared by ESPript 3.0 (https://espript.ibcp.fr/ESPript/cgi-bin/ESPript.cgi). The black triangle indicates CvkR lacks of a cysteine residue in the C-terminal α-helical domain which is conserved in the selected members of the MerR-type regulators.

### Supplemental Tables

**Table S1. Details of 118 detected CAST systems in cyanobacteria.**

Accession numbers and deduced amino acid sequences are given for all identified Cas12k and CvkR homologs (columns A to F), followed by the respective strain names and genome or contig accession numbers (columns G and H). The insertion elements LE always refer to the IS-element closer to the *cas12k* gene and RE refers to the opposite end (columns I and J). The detected *cas12k* and *cvkR* genes were compared to each other and in some cases, we corrected the start codon positions compared to the NCBI annotation, as detailed for *all3613* and *alr3614* in **Fig. 3** of this work. Those instances of proteins with corrected start codon positions are labeled by the [§] suffix in columns A and D and the resulting shorter sequences are given in columns C and F. Some CAST systems showed likely pseudogenized *cas12k* genes indicated by the suffix [#] in column A. If multiple CAST systems exist in one strain, these were numbered consecutively in column G.

**Table S1** is provided as separate Excel file.

**Table S2. Microarray analysis of Δ*cvkR* or Δ*cvkR*Com deletion and complementation mutants.** Microarray analysis result table of significantly changed features in Δ*cvkR* compared to the complementation strain Δ*cvkR*Com. Threshold: $|\log_2| \geq 1$, p-value ≥0.01. Two replicates were used for the microarray experiment.

| **Probe_Label** | **Gene/region_Function** | **location** | **log$_2$FC** | **adj. pvalue** |
|---|---|---|---|---|
| alr0738_igFwd | NA | chr | -2.413 | 1.41E-03 |
| *rtcB* | RNA-splicing ligase | chr | -2.382 | 4.19E-03 |
| *cvkR* | CvkR | chr | -2.203 | 4.77E-04 |
| nTSS_52243_52243 | Upstream of tRNA | delta | -1.987 | 4.77E-04 |
| *all8564* | Restriction endonuclease (5-methylcytosine-specific) | delta | -1.743 | 1.41E-03 |
| *alr0739* | Uncharacterized conserved protein YdeI | chr | -1.616 | 5.37E-04 |
| *alr0740* | slipin family protein | chr | -1.500 | 1.06E-04 |
| *asr0855* | Unknown | chr | -1.328 | 4.19E-03 |
| *all7121* | Cytochrome c domain-containing protein | alpha | 1.075 | 1.41E-03 |
| all0328_igFwd | NA | chr | 1.109 | 3.81E-03 |
| alr0786_igRev | NA | chr | 1.268 | 3.28E-03 |
| *pecB* | phycoerythrocyanin Beta-chain | chr | 1.300 | 4.19E-03 |
| all0736_igRev | NA | chr | 1.344 | 8.26E-03 |
| alr1198_igFwd | NA | chr | 1.387 | 1.12E-03 |
| *pecA* | phycoerythrocyanin Alpha-chain | chr | 1.416 | 1.68E-03 |
| *pecC* | phycoerythrocyanin-associated rod linker protein | chr | 1.435 | 1.69E-04 |
| tracrRNA | CAST tracrRNA | chr | 1.493 | 4.33E-03 |
| *cas12k* | CAST effector gene | chr | 1.550 | 1.41E-03 |
| all3391_igFwd | NA | chr | 1.567 | 4.77E-04 |
| *tnsB* | CAST transposase | chr | 1.594 | 4.67E-04 |
| *cas12k* | CAST effector gene | chr | 1.870 | 9.16E-05 |

**Table S3. Details of crystallographic analysis.**

| | SeMet CvkR | Native CvkR |
|---|---|---|
| **Data collection** | | |
| Space group | C121 | C121 |
| Unit cell | | |
| a, b, c (Å) | 78.813, 49.778, 41.017 | 78.597, 49.675, 40.506 |
| α, β, γ (°) | 90, 116.38, 90 | 90, 116.055, 90 |
| Wavelength (Å) | 0.97918 | 0.97930 |
| Resolution (Å)[a] | 24.90-1.93 (2.03-1.93) | 50.00-1.50 (1.53-1.50) |
| Total reflections | 33044 | 132616 |
| Unique reflections | 10215 | 22409 |
| $I/\sigma(I)$ | 14.6 (3.8) | 42.57 (1.57) |
| Completeness (%) | 94 (84.4) | 99.1 (89.7) |
| Redundancy | 3.2 (3.1) | 5.9 (4.2) |
| $R_{merge}$ (%) | 5.4 (25) | 4.1 (51.4) |
| **Phasing** | | |
| Figure of merit[b] | 0.55 | |
| BAYES-CC | 0.33 | |
| Sites | 11 | |
| **Refinement** | | |
| $R_{work}/R_{free}$ (%) | | 18.81/22.66 |
| **B-factors** | | 46. 42 |
| Protein | | 45. 71 |
| ATP | | 60.14 |
| water | | 52.51 |
| **r.m.s.[c] deviations** | | |
| Bond lengths (Å) | | 0.01 |
| Bond angles (°) | | 1.30 |
| **Ramachandran Plot (%)** | | |
| Favored | | 99 |
| Allowed | | 0.68 |
| Outliers | | 0 |

[a] The highest resolution shell is shown in parentheses.
[b] Probability weighted average of the cosine of the phase error before and after density modification.
[c] Root mean square.

**Table S4. Oligonucleotide primers used in this work.** Sequences belonging to the T7 promoter are underlined. Primers were purchased from IDT or Tsingke.

| | | |
|---|---|---|
| *alr3614* deletion mutant (Δ*cvkR*) construction | | |
| alr3614gRNA-1 | AGATCATTAACAGTAATGGAGCAG | gRNA for *alr3614* |
| alr3614gRNA-2 | AGACCTGCTCCATTACTGTTAATG | |
| alr3614KO-1 | GGTCATTTTTTTGTCTAGCTTTAATGCGGTAGTTGGTACCGGTTAAGAGAATATCCTGCC | 5' homology arm for *alr3614* knock out |
| alr3614KO-2 | CAGGCGATCGCGTGGGTAATTACTATAACTCCTTTCTCTC | |
| alr3614KO-3 | GAGAGAAAGGAGTTATAGTAATTACCCACGCGATCGCCTG | 3' homology arm for *alr3614* knock out |
| alr3614KO-4 | GCGCTGCCCGGATTACAGATCCTCTAGAGTCGACGGTACCAAAATCTTCGATGCAGATGA | |
| alr3614-3 | GACGACTATCGCAGTAAACTTC | Genotype confirmation |
| alr3614-4 | TTAGCTGCTTTACTACGACG | |
| *alr3614* complementation strain (Δ*cvkR*Com) construction | | |
| 59M-F | GATTATAAAGATCATGATGGTGATTATAAAGATCATGATATTGATTATAAAGATGATGATGATAAATGAAAGGGTGGGCGCGCCGACCCAG | Amplification of pRL59EH-Cm/Em backbone with overlap to 3xFlag tag |
| 59M-R1 | GTTAATTTCACAGGCTTTAGGTCGACCTGCATCCCTTAAC | Amplification of pRL59EH-Cm/Em backbone with overlap to $P_{petE}$ |
| $P_{petE}$-F | GTTAAGGGATGCAGGTCGACCTAAAGCCTGTGAAATTAAC | With overlap to pRL59EH-Cm/Em backbone |
| $P_{petE}$-R4 | CACATTAGTGCTTTGTATAACATTTATTTCATTTTAAATAAAATCGACACC | $P_{petE\_no5'UTR}$-*alr3614*L-3xFlag |
| $P_{petE}$-R5 | GCTTGTGTAGAACTTTCCTTCCATTTATTTCATTTTAAATAAAATCGACACC | $P_{petE\_no5'UTR}$-*alr3614*S-3xFlag |
| $P_{petE}$-R6 | GTGTAGAACTTTCCTTCCATGGCGTTCTCCTAACCTGTAG | $P_{petE}$-*alr3614*S-3xFlag |
| 3614L-F | GGTGTCGATTTTATTTAAAATGAAATAAATGTTATACAAAGCACTAATGTG | $P_{petE\_no5'UTR}$-*alr3614*L-3xFlag |
| 3614S-F1 | CGATTTTATTTAAAATGAAATAAATGGAAGGAAAGTTCTACACAAGC | $P_{petE\_no5'UTR}$-*alr3614*S-3xFlag |
| 3614S-F2 | CTACAGGTTAGGAGAACGCCATGGAAGGAAAGTTCTACAC | $P_{petE}$-*alr3614*S-3xFlag |
| 3614-R | TCATTTATCATCATCATCTTTATAATCAATATCATGATCTTTATAATCACCATCATGATCTTTATAATCGCTACTAAAGCTTTTAAGATTC | Amplication of *alr3614*-3xFlag with overlap to pRL59EH-Cm/Em backbone |
| 3614TesT-F | GCCGCCAGTTGCAGTATT | Genotype confirmation |
| 3614TesT-R | CGGGCAAGTACGACATCA | |
| qRT-PCR | | |
| all3613 RT-F | TCTATCCCTTTTCCTGTGGT | |
| all3613 RT-R | CGTTTAGTTTGTTGGTCTTCC | |
| tracrRNA RT-F | TAAGGTTTTCAGGATGTGCG | |
| tracrRNA RT-R | TGCCAATAGACAGGATAGGT | |
| pre-crRNA RT-F | GATCGCGCACAATATAAAG | |
| pre-crRNA RT-R | GCTGATGAAGAAAAAGTTAG | |
| alr3614 RT-F | GTGGCAAAGGTCGTAATGTT | |
| alr3614 RT-R | CGCTTCATCTTTTCTTCTGGG | |
| rnpB RT-F | CGTGAGGATAGTGCCACAGA | |

| | | |
|---|---|---|
| rnpB RT-R | CCAACCATAGTTCCTTCGGC | |
| **Northern hybridizations** | | |
| CR_9_fwd | TTTGAATATTCAGAACTTTATATTGTGCGCGAT | as published[3], for mutant analysis |
| CR_9_rev | TAATACGACTCACTATAGGGGCAAGCTGATTTGGTAGAAGCTGTTAAT | for mutant analysis |
| all3613-Nblot1 | GGAGTAAGCTTGGGGCTAGAA | for mutant analysis |
| all3613-Nblot2 | TAATACGACTCACTATAGGGGCCTGCTTTATAGGTCTGTGC | |
| tracrRNA_Fw_T7_1 | TAATACGACTCACTATAGGGCTCTTTGGTGCGTCAAATCAAG | for mutant analysis |
| tracrRNA_Rev | CAGTTCATGCTGCTTGCAGC | |
| Alr3614_Nblot_f | GCACAGAAGCATCAGAAATTAC | for mutant analysis |
| Alr3614_Nblot_r | TAATACGACTCACTATAGGGAGACAGCAACTGCTCCATTAC | |
| 5S_7120 | TAGCAGCGTTTCACCTCTGAGTTCGG | as published[3] |
| **TXTL assay** | | |
| pet28a_HisTEV_fwd | CTCGAGCACCACCACCAC | Amplification of pET-28a(+) with 6xHis tag and TEV site |
| pet28a_HisTEV_fwd | GGATCCCATATGCTGAAAATACAGG | |
| 3614mut_fwd | ATTTTCAGCATATGGGATCCATGGAAGGAAAGTTCTACAC | Amplification of *cvkR*mut with overlaps to pET-28a(+) |
| 3614mut_rev | TGGTGGTGGTGGTGCTCGAGTTAGCTACTAAAGCTTTTAAGATTC | |
| p70a-deGFP_fwd | GCTAGCAATAATTTTGTTTAACTTTAAGAAGGAGATATACC | Amplification of p70a with deGFP and 5´UTR but without promoter |
| p70a-deGFP_rev | GCATGCCCAGCGGAACAG | |
| P3613_fwd | TGCTGTTCCGCTGGGCATGCAAGAAACATCCTATAGAAGC | Amplification of the *all3613* promoter with overlaps to p70a |
| P3613_rev | TAAACAAAATTATTGCTAGCAAAAATAAAATACCATACAAAACAC | |
| P3614_fwd | TGCTGTTCCGCTGGGCATGCAAAAATAAAATACCATACAAAACAC | Amplification of the *alr3614* promoter with overlaps to p70a |
| P3614_rev | TAAACAAAATTATTGCTAGCAAGAAACATCCTATAGAAGC | |
| P43_fwd | TGCTTTGTATAACATTATGTGTTTTGTATGGTATTTTATTTTTGCTAGCAATAATTTTGTTTAACTTTAAGAAGGAGATATACC | Amplification of p70a with deGFP under control of a shorter version of the *all3613* promoter |
| P43_rev | AAAAATAAAATACCATACAAAACACATAATGTTATACAAAGCAGCATGCCCAGCGGAACAG | |
| P39_fwd | TAAAACACATTAGTGCTTTGTATAACATTATGTGTTTTGGCTAGCAATAATTTTGTTTAACTTTAAGAAGGAGATATACC | Amplification of p70a with deGFP under control of a shorter version of the *all3613* promoter |
| P39_rev | CAAAACACATAATGTTATACAAAGCACTAATGTGTTTTAGCATGCCCAGCGGAACAG | |
| P26_fwd | TGCTTTGTATAACATTATGTGTTTTGGCTAGCAATAATTTTGTTTAACTTTAAGAAGGAGATATACC | Amplification of p70a with deGFP under control of a shorter version of the *all3613* promoter |
| P26_rev | CAAAACACATAATGTTATACAAAGCAGCATGCCCAGCGGAACAG | |
| P20_fwd | GTATAACATTATGTGTTTTGGCTAGCAATAATTTTGTTTAACTTTAAGAAGGAGATATACC | Amplification of p70a with deGFP under control of a shorter version of the *all3613* promoter |
| P20_rev | CAAAACACATAATGTTATACGCATGCCCAGCGGAACAG | |
| Ptracr_fwd | TGCTGTTCCGCTGGGCATGCCGCAGGATAAAGCAAAAG | |

| | | |
|---|---|---|
| Ptracr_rev | TAAACAAAATTATTGCTAGCTATTACATATTATAT TTTCAAGGTTCAG | Amplification of p70a with deGFP under control of the tracrRNA promoter |
| Alr3614S protein heterogenous expression | | |
| 3614S-BamHI-F | CGGGATCCATGTTATACAAAGCACTAATGTG | Producing 3614S-pET-28a(+)-smt3 |
| 3614S-taa-XhoI-R | CCGCTCGAGTTAGCTACTAAAGCTTTTAAGATTC TC | |
| Promoters for EMSA and/or TXTL assays (Blue: -10 element; Green: -35 element) | | |
| $P_{cas12k}$=P82 | AAGAAACATCCTATAGAAGCATCTTCTAAAACACA TTAGTGCTTTGTATAACATTATGTGTTTTGTATGGT ATTTTATTTTT | |
| $P_{tracr}$ | CGCAGGATAAAGCAAAAGAATTAGCCCTCTATGC TTATAGTCTCCGCCTAGCTAGGCGAAGTTGACAA ACATCTGAACCTTGAAAATATAATATGTAATA | |
| $P_{cvkR}$ | AAAAATAAAATACCATACAAAACACATAATGTTATA CAAAGCACTAATGTGTTTTAGAAGATGCTTCTATA GGATGTTTCTT | |
| P43 | TGCTTTGTATAACATTATGTGTTTTGTATGGTATTT TATTTTT | |
| P39 | TAAAACACATTAGTGCTTTGTATAACATTATGTGTT TTG | |
| P26 | TGCTTTGTATAACATTATGTGTTTTG | |
| P20 | GTATAACATTATGTGTTTTG | |
| DNase I footprinting | | |
| P3613-F(FAM) | AACTCCTTTCTCTCGCCAATAC | Producing FAM-labeled probes |
| P3613-R | GCTGCTGAAGCAGTTCGTT | |
